# Her9 controls the stemness properties of the hindbrain boundary cells

**DOI:** 10.1101/2024.03.12.584657

**Authors:** Carolyn Engel-Pizcueta, Covadonga F Hevia, Adrià Voltes, Jean Livet, Cristina Pujades

## Abstract

Different spatiotemporal distribution of progenitor/neurogenic capacities permits that brain regions engage asynchronously in neurogenesis. In the hindbrain, rhombomere progenitor cells are the main contributors to neurons during the first neurogenic phase, whereas boundary cells participate later, relying on Notch3-activity. To analyze the mechanism(s) that maintain boundary cells as proliferative progenitors not engaging in neurogenesis, we addressed the role of the zebrafish Hes1 ortholog, Her9, in this cell population. *her9* expression is temporarily sustained in boundary cells in a Notch-independent manner while they behave as non-neurogenic progenitors. Functional manipulations demonstrate that Her9 inhibits the onset of Notch-signaling and the neurogenic program, thus keeping boundary cells in the progenitor state. Combining multicolor clonal analysis with functional approaches, we reveal a role of Her9 in the expansion of boundary progenitors by promoting symmetric proliferative divisions and preventing neurogenic cell divisions. Moreover, Her9 regulates the proliferation of boundary cells by inhibiting the cell cycle gene *cdkn1ca* and potentially interplaying with CyclinD1. Altogether, Her9 maintains the stemness and proliferation of hindbrain boundary progenitors at early embryonic stages.

## INTRODUCTION

During early brain development, neuroepithelial cells, a primary form of neural stem cells, proliferate by symmetric cell divisions to expand the population and contribute to the growth of the neural tube. Subsequently, neuroepithelial cells transition into radial glial cells, which are neural progenitors that divide asymmetrically, giving rise to neurons while maintaining the stem cell pool. Timely shifts along cell division modes are crucial to form a functional brain. In the hindbrain, the embryonic brainstem, the orchestrated spatiotemporal distribution of progenitor and neurogenic capacities results in hindbrain territories engaging asynchronously in neurogenesis (Nikolaou *et al*, 2009; Hevia *et al*, 2022; Voltes *et al*, 2019; Peretz *et al*, 2016; Gonzalez-Quevedo *et al*, 2010; Belmonte-Mateos *et al*, 2023). This asynchrony relies on a spatial organization that results from hindbrain segmentation and gives rise to seven transient rhombomeres (Kiecker & Lumsden, 2005). At the interface between rhombomeres, there is the specification of the boundary cell population (Guthrie & Lumsden, 1991; Lumsden & Keynes, 1989), which exhibits specific gene expression (Cheng *et al*, 2004; Letelier *et al*, 2018; Moens *et al*, 1996) and displays distinct functions during development (Pujades, 2020). Rhombomeres engage very early in neurogenesis (Nikolaou *et al*, 2009; Gonzalez-Quevedo *et al*, 2010; Cheng *et al*, 2004; Amoyel *et al*, 2005), whereas boundary cells participate later (Hevia *et al*, 2022; Voltes *et al*, 2019; Peretz *et al*, 2016). This distinct neurogenic commitment is mainly regulated by Notch-signaling. During the early neurogenic phase, Notch-pathway is highly active in the neurogenic rhombomere compartments (Nikolaou *et al*, 2009), while inactive in the hindbrain boundaries composed by neuroepithelial cells dividing in a proliferative symmetric mode (Hevia *et al*, 2022; Voltes *et al*, 2019). Later, Notch3-activity ensues the switch of boundary cells to radial glia progenitors that divide asymmetrically (Hevia *et al*, 2022). This is a common scenario in other neural systems, in which neurogenic commitment arises through Notch-signaling conferring the capacity to progenitors to divide asymmetrically (Nerli *et al*, 2020; Mizutani *et al*, 2007; Kressmann *et al*, 2015; Dong *et al*, 2012). We unveiled that Yap/Taz-activity maintains boundary cells as highly proliferative progenitors (Voltes *et al*, 2019). However, the mechanism(s) preventing boundary cells from transitioning to neurogenesis and the cell cycle regulators controlling their proliferative capacity at early embryonic stages are still unknown.

The Hes family of basic Helix-Loop-Helix (bHLH) transcriptional repressors is key in the balance between cell differentiation and proliferation by maintaining the neural stem cells through the inhibition of proneural transcription factors and cell cycle exit genes (Kageyama *et al*, 2008; Ohtsuka & Kageyama, 2021). Although Hes factors are the main effectors of Notch-signaling in neurogenesis (Ohtsuka *et al*, 1999), there is a set of Hes genes expressed in high and sustained levels which expression is independent of Notch signaling (Cau *et al*, 2000; Geling *et al*, 2003; Ninkovic *et al*, 2004). In teleosts, Hes/Her maintain neural progenitors at the mid-hindbrain boundary by inhibiting neurogenesis in a Notch-independent manner (Geling *et al*, 2003; Ninkovic *et al*, 2004). In mammals, Hes1 is a key factor for maintaining neural stem cells in the undifferentiated state. Brain boundaries express high and constant Hes1, whereas compartments show an oscillatory Hes1 expression (Baek *et al*, 2006; Shimojo *et al*, 2008). Loss of *Hes1* results in ectopic proneural gene expression and premature neuronal differentiation (Cau *et al*, 2000; Hirata *et al*, 2001; Ohtsuka *et al*, 1999; Ishibashi *et al*, 1995; Hatakeyama *et al*, 2004), whereas high *Hes1* expression represses neurogenesis and keeps neural stem cells by maintaining symmetric proliferative divisions (Sueda *et al*, 2019; Ohtsuka & Kageyama, 2021). Accordingly, sustained Hes1 expression in neural progenitors leads to inhibition of proneural genes, delta ligands, Notch effectors (Hes5), and cell cycle-related genes such as *Ccnd1* (Baek *et al*, 2006; Shimojo *et al*, 2008; Sueda *et al*, 2019; Ohtsuka & Kageyama, 2021).

To further analyze the mechanism(s) that maintain cells as proliferative progenitors before undergoing neurogenesis, we addressed the role of the zebrafish ortholog of Hes1, Her9, in hindbrain boundary cells. We demonstrate that *her9* is enriched in a Notch-independent manner in boundary cells when they are behaving as non-neurogenic progenitors. Manipulation of Her9 reveals that it represses the onset of Notch-signaling and the neurogenic program, thus keeping boundary cells in the progenitor state. Functional multicolor clonal analyses show that Her9 expands the boundary progenitor population by promoting symmetric proliferative and preventing neurogenic cell divisions. Furthermore, Her9 inhibits the cell cycle gene *cdkn1ca* and potentially cooperates with CyclinD1 to control the proliferative capacity of boundary cells.

## RESULTS

### *her9* is enriched in the hindbrain boundaries independently of Notch signaling

To explore the role of the zebrafish ortholog of Hes1, Her9, on hindbrain boundary cells we first analyzed the expression of *her9* in the hindbrain. We performed double *in situ* hybridizations with *her9* and well described hindbrain boundary markers such as *rfng* and *foxb1a* (Cheng *et al*, 2004; Moens & Prince, 2002). At early embryonic stages, *her9* was enriched in the hindbrain boundaries where it largely overlapped with boundary markers (Figure 1A–A’’, a–a’’; Figure EV1A–A’’, a–a’’). Next, we compared its expression to *her4*, a classical Notch effector expressed in neurogenic progenitors (Stigloher et al., 2008; Takke et al., 1999). We observed *her4* was expressed all along the rhombic lip, although its expression was restricted to rhombomere compartments and devoid in the boundaries (Figure 1B–B’, b– b’). Thus, at the stage when hindbrain boundaries are not neurogenic *her9* and *her4* did not overlap (Figure 1B’’, b’’), showing a complementary expression. Spatiotemporal analysis of *her9* expression in the hindbrain revealed that high *her9* in the boundaries started at 18hpf (Figure 1C, c), coinciding with the onset of expression of boundary specific genes (Letelier *et al*, 2018). *her9* expression in the progenitor domain of the hindbrain boundaries was sustained upon lumen formation (18-30hpf; Figure 1C–E, c–e), whereas *her9* showed lower levels within rhombomeres (Figure 1C–E). From 26hpf onwards, *her9* became enriched in two lateral stripes all along the anteroposterior (AP) axis (Figure 1E–F, e–f; Figure EV1A, C, a, c). From 32hpf onwards, the enriched *her9* expression in the boundaries was lost and the expression became homogenous in boundary and rhombomere compartments (Figure 1F, f; Figure EV1C, c). Consistent with this, by 36hpf most boundary cells expressed the Notch effector *her4* (Figure EV1B–B’, b–b’), and only a few still expressed *her9* (Figure EV1C, c). Hence, the decrease of *her9* in boundary cells coincides with the onset of Notch signaling and thus, the expression of *her4* and the engagement in neurogenesis (Hevia *et al*, 2022). Overall, *her9* is dynamically expressed in distinct progenitor domains during early embryonic stages of hindbrain development.

**Figure 1.**
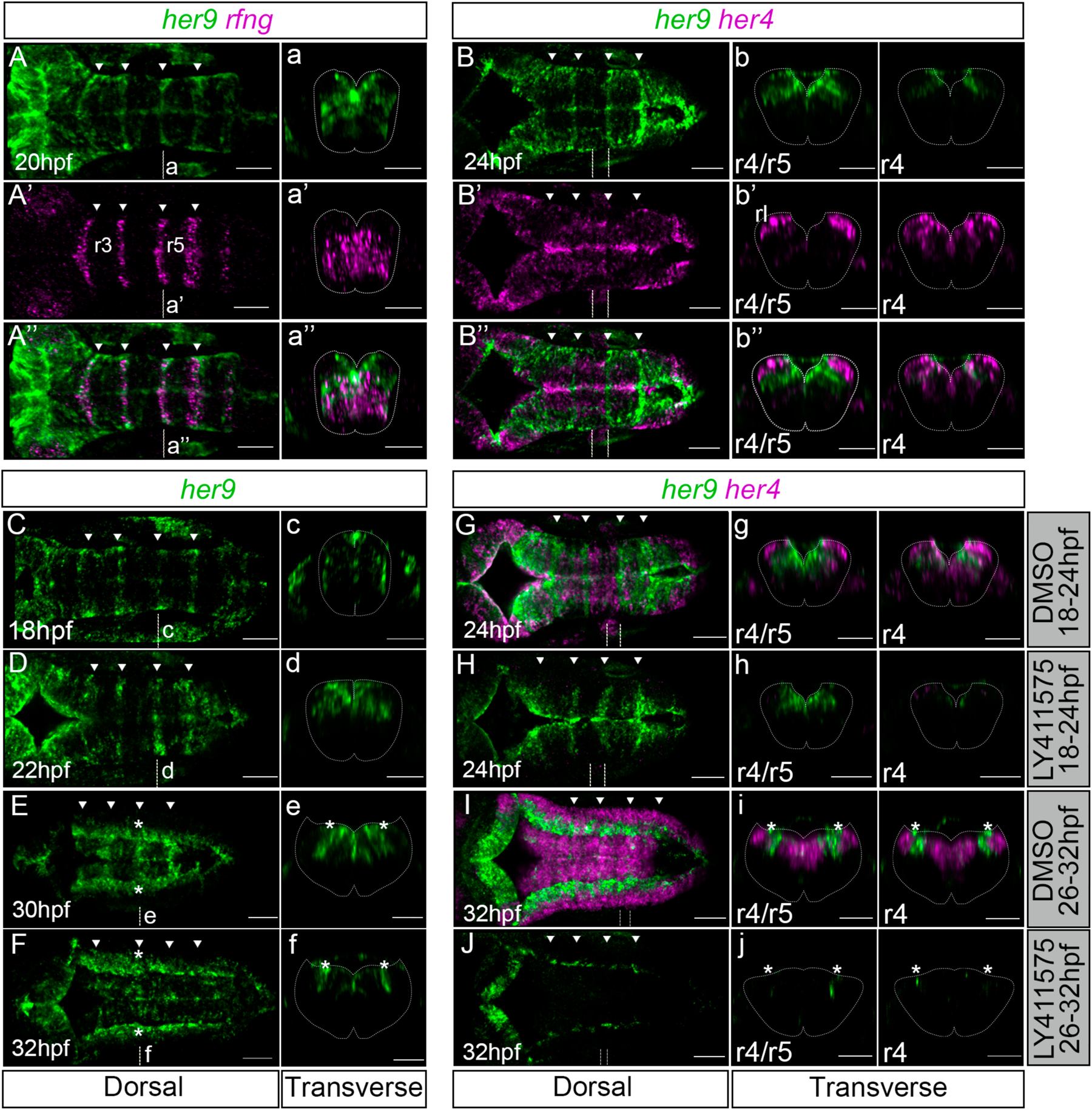
*her9* is enriched in the hindbrain boundaries in a Notch independent manner at early embryonic stages. (A–J) *her9 in situ* hybridization with *rfng* (A’–A’’) and *her4* (B’–B’’, G–J) at the indicated developmental stages. (G–J) Embryos treated with DMSO (G, I) or LY411575 (H, J) at the two indicated stages (18-24hpf: n=4/4 DMSO vs. n=14/14 LY411575; 26-32hpf: DMSO n=10/10 vs. LY411575 n=8/9). (A–A’, B–B’, C–F) Single and (A’’–B’’, G–J) merged channels. (A–A’’, B–B’’, C–J) Dorsal MIP (Maximal Intensity Projection) of the hindbrain with anterior to the left, with (a–a’’, b–b’’, c–j) transverse views of r4/r5 boundary (if not indicated) or r4. White arrowheads indicate the position of the hindbrain boundaries. Dashed white line delimitates the contour of the neural tube. hpf, hours post fertilization, r, rhombomere; rl, rhombic lip. Scale bar 50μm.

To explore whether sustained *her9* expression was dependent on Notch signaling, we downregulated Notch activity using the gamma-secretase inhibitor LY411575. We targeted two different time intervals: i) 18 to 24hpf, when *her9* was highly enriched in hindbrain boundaries, and ii) 26 to 32hpf, when *her9* expression in the boundaries decreased. In the first case, upon Notch downregulation sustained *her9* expression was maintained in the hindbrain boundaries although it was inhibited in rhombomeres when compared to controls (Figure 1G–H, g–h); accordingly, boundary identity was not compromised (Figure EV1D–E, d– e). Consistent with *her4* being a Notch target, its expression was decreased upon Notch downregulation (Figure G–H, g–h). On the other hand, when Notch was downregulated at later stages, both *her4* and *her9* expression greatly decreased when compared to controls, except for some remaining *her9* in the most lateral domains (Figure 1I–J, i–j). This coincided with the ceasing of expression of boundary cell markers (Figure EV1F, f; (Letelier *et al*, 2018). Therefore, the sustained expression of *her9* in the hindbrain boundaries is Notch-independent, consistent with the absence of Notch activity in this structure at this stage (Hevia et al, 2022).

### Her9 maintains the stemness of hindbrain boundary cells at early embryonic stages

To determine whether Her9 could maintain boundary cells in the progenitor state, we knocked down *her9* by injecting one-cell stage embryos with a splice-blocking morpholino (her9-MO; Figure EV2A) and analyzed its impact on boundary cell fate. First, we evaluated the efficiency of the her9-MO by assessing the defects in the opening of the fourth ventricle described by (Bae *et al*, 2005) using a cadherin reporter line (Revenu *et al*, 2014). her9-MO embryos showed an aberrant opening of the ventricle and the neural tube with a rounded shape in contrast to controls (Figure EV2B–C, b–c). This phenotype correlated with the presence of defective *her9* spliced forms (Figure EV2D). Then, to study the role of Her9 in boundary cells, we injected embryos, in which differentiated neurons and boundary cell nuclei were labeled, with control-MO and her9-MO. When almost no boundary cells engage in neurogenesis (30hpf), we observed an increase in the percentage of boundary-derived neurons upon *her9* downregulation compared to control-MO embryos (Figure 2A–C, A’–B’, a’–b’). However, at 36hpf, when boundary cells had started the neurogenic program, no significant differences in boundary derived neurons were observed (Figure 2D–F, D’–E’, d’– e’). These results suggest that Her9 keeps boundary cells as progenitors preventing them to engage in neurogenesis at early embryonic stages. When the effect of Her9 on neuronal commitment genes such as *neuroD4* was assessed, an increase in the percentage of *neuroD4*-boundary cells in her9-MO was observed (Figure 2G–I, G’–H’, g’–h’). Therefore, *her9* downregulation causes a premature engagement of boundary cells into neurogenesis.

**Figure 2.**
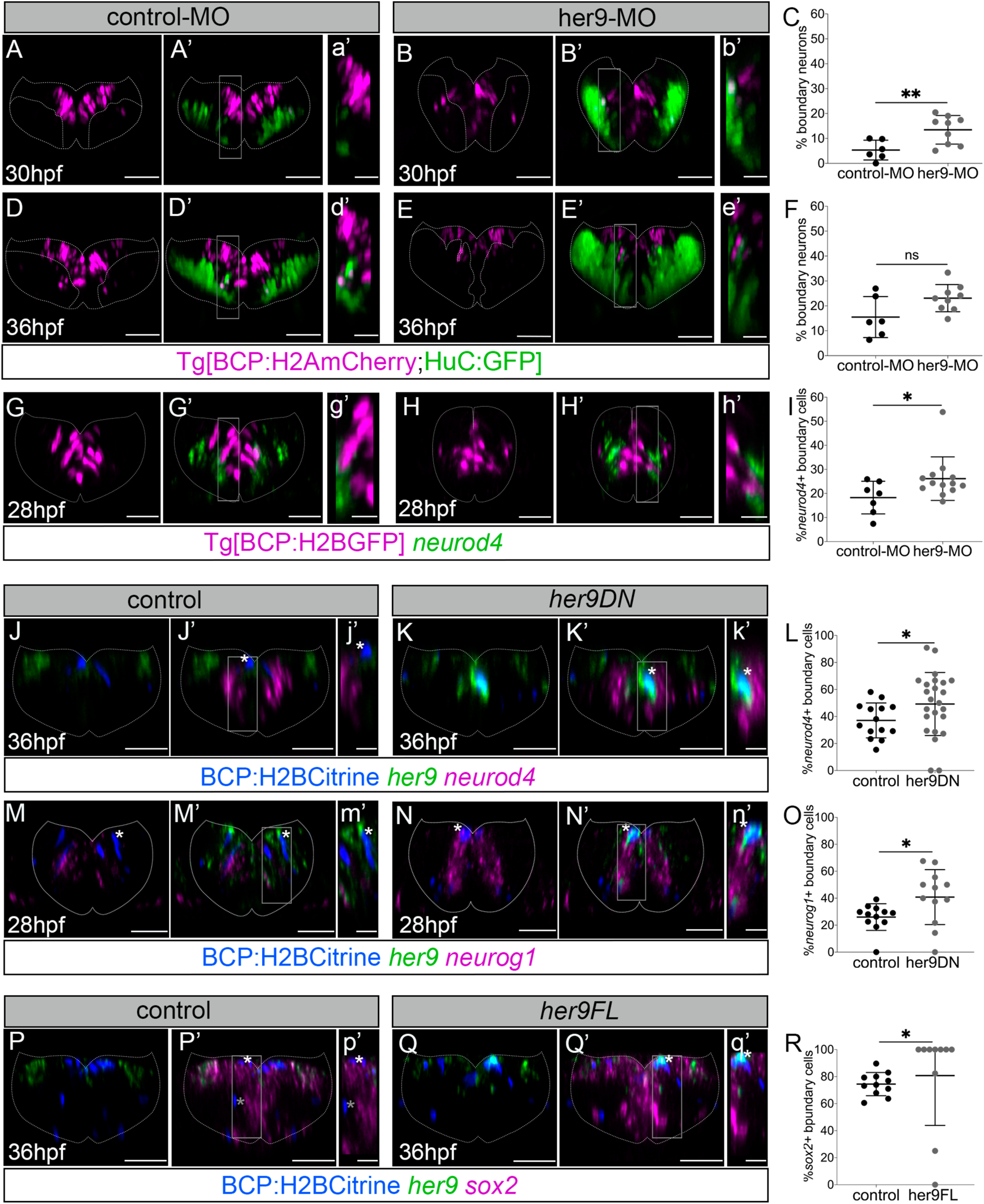
Her9 maintains boundary cells as progenitors preventing them to engage in neurogenesis. (A–B, D–E, A’–B’, D’–E’) Tg[BCP:H2AmCherry;HuC:GFP] embryos with boundary cells nuclei in magenta and differentiated neurons in green injected with control-MO (A–A’, D–D’) or her9-MO (B–B’, E–E’) and analyzed at the indicated times. (C, F) Plots displaying the percentage of boundary derived neurons at 30hpf and 36hpf (30hpf: control-MO 5.3% ± 4 n=6 vs. her9-MO 13.5% ± 5.7 n=9, p value=0.007 [**], Welch’s test; 36hpf: control-MO 15.5% ± 8.2 n=6 vs. her9-MO 23.1% ± 5.5 n=9, p value=0.08 [ns], Welch’s test). (G–H, G’–H’) Tg[BCP:H2BGFP] embryos injected with control-MO (G–G’) or her9-MO (H–H’) and *in situ* hybridized with *neuroD4* at 28hpf. (I) Plot showing the percentage of boundary cells expressing *neurod4* (control-MO 18.2% ± 6.8 n=7 vs. her9-MO 26.1% ± 9.1 n=13; p value=0.047 [*], Mann-Whitney test). (J–Q, J’–Q’) Tg[BCP:Gal4] embryos injected with H2Bcitrine:UAS (J–J’, M–M’, P–P’), H2Bcitrine:UAS:her9DN (K–K’, N–N’) or H2Bcitrine:UAS:her9FL (Q–Q’) and *in situ* hybridized with *her9* and *neurod4* (J–K, J’–K’), *neurog1* (M–N, M’–N’), or *sox2* (P–Q, P’–Q’). Note that control H2Bcitrine-boundary cells (blue) do not express *neurod4* or *neurog1* (j’, m’; see asterisks), whereas H2Bcitrine:her9DN-cells colocalized with *neurod4* or *neurog1* (k’, n’; see asterisks). Boundary cells in the ventricular domain express *sox2* and those in the ventral domain do not (P’, p’; see white and gray asterisks) whereas H2Bcitrine:her9FL-cells express both *her9* and *sox2* (Q’, q’; see white asterisk). (L, O, R) Plots with percentage of boundary cells expressing *neurod4* (control: 37.1% ± 12.9 n=14 boundaries, N=4 embryos vs. her9DN: 49.2% ± 23.4 n=26 boundaries, N=7 embryos; p value=0.046 [*], Welch’s test), *neurog1* (control: 26.1% ± 9.8 n=12 boundaries, N=6 embryos vs. her9DN: 40.8% ± 20.4 n=12 boundaries, N=7 embryos]; p value=0.038 [*], Welch’s test), or *sox2* (control: 74.5% ± 8.5 n=11 boundaries, N=4 embryos vs. her9FL: 80.7% ± 36.9 n=12 boundaries, N=5 embryos; p value=0.026 [*], Mann-Whitney test). All plots show the mean ± SD. Transverse views of r4/r5 except for (P–Q, P’–Q’) that correspond to r3/r4 boundary. (a’–q’) Magnifications of the framed regions in (A’–Q’). Dashed white line delimitates the contour of the neural tube. BCP, Boundary Cell Population; hpf, hours post fertilization; MO, morpholino. Scale bar 50μm, and 20μm for magnifications.

To specifically manipulate Her9 in boundary cells, we performed conditional Her9 loss-and gain-of-function assays and analyze the impact on boundary cell fate. We generated embryos with boundary cells expressing either a dominant negative form of *her9* (her9DN) or the full-length *her9* (her9FL) by injecting UAS-driven her9DN/FL constructs in Tg[BCP:Gal4] embryos (Figure EV2E). First, we assessed the neuronal fate of boundary cells expressing the her9DN at 36hpf. Half of these cells expressed *neuroD4,* a significant increase compared to control cells (Figure 2J–L, J’–K’, j’–k’). To confirm this was due to an increase in neuronal specification, we analyzed the expression of *neurog1* at 28hpf, one of the first proneural genes to act in this process (Guillemot, 2007). The percentage of her9DN-positive boundary cells expressing *neurog1* was higher when compared to control cells (Figure 2M–O, M’–N’, m’–n’). Thus, sequestering Her9’s function increased the number of boundary cells committed to the neuronal lineage. To further investigate how Her9 prevented boundary cells from entering neurogenesis, we monitored Notch activity upon *her9* downregulation. We found an increase of Notch-active boundary cells already at 30hpf in her9-MO embryos compared to control-MO ones (Figure EV3A–C, A’–B’, a’–b’). We further detected a significant increase of her9DN cells expressing *deltaD*, the main Notch ligand expressed in the boundaries at 28hpf (Hevia *et al*, 2022) in contrast to control cells (Figure EV3D–F, D’–E’, d’–e’). These results indicate that Her9 may inhibit *deltaD* expression, and thus Notch activity, preventing the onset of neurogenesis in the hindbrain boundaries. Next, we determined whether Her9 was also sufficient to maintain boundary cells as progenitors. When we conditionally overexpressed her9FL in the boundaries we observed an increase of these cells in the progenitor *sox2* domain when compared to control cells (Figure 2P–R, P’–Q’, p’–q’). Hence, maintaining sustained *her9* expression keeps boundary cells in the progenitor state. Altogether, these data indicate that Her9 is necessary and sufficient to maintain the stemness of boundary cells by inhibiting the expression neurogenic genes.

### Her9 controls the behavior of boundary cells

Hindbrain boundary cells transition from symmetrically dividing progenitors to asymmetrically dividing cells relying on Notch3 activity (Hevia *et al*, 2022), coinciding with the loss of *her9* enrichment. Since we wanted to unveil the role of Her9 in boundary cells, we first established a system to better understand how this transition occurs at the single cell level, to be able later to assess the phenotype upon Her9 disruption. Thus, we performed multicolor clonal analysis taking advantage of the zebrabow1.0 system (Pan *et al*, 2013), which enables to randomly label cells in different colors in such a manner that clonally related cells display the same color. Thus, we generated colored boundary clones and tracked them *in vivo* from 32 (t0) to 45hpf (tf) to reconstruct their lineage while assessing their mode of cell division (Video 1; Figure 3A–C, b–c). We combined color and spatial criteria to simultaneously ascribe lineage and fate (Figure EV4). During the analyzed temporal window, half of the progenitors divided, and the other half did not divide (Figure 3D). At tf, one third of the tracked boundary cells had undergone neurogenesis while the rest remained as progenitors (Figure 3E), consistent with our previous results on boundary cell lineage (Hevia *et al*, 2022). Most clones had two cells at t0 (n=34/44) and at tf, they were composed of two (n=16/44) or four cells (n=17/44; Figure 3F). The most frequent cases being that of two-cell clones in which no cell divided (n=12/34; Figure 3F, t0) or of two-cell clones in which both cells divided (n=13/34; Figure 3F, tf). Upon analysis of the cell division mode, we observed that progenitor cells mainly divided proliferative symmetrically (PP) or asymmetrically (PN), with only one single case of neurogenic symmetric division (NN) detected (Figure 3G). No temporal distribution of PP and PN divisions in boundary cells was observed (Figure 3H) in accordance with (Hevia *et al*, 2022). Therefore, no preference in the cell division mode (PP or PN) was found in terms of the number of divisions that the clone underwent or the time that cells divided. Sister cells tended to have the same proliferative capacity, namely either both cells in the clone divided or they did not (Figure 3I), although they could undergo the same or a different cell division mode (Figure 3J). Thus, both sister cells could display PP or PN divisions (Figure 3K as an example of PN; top panel) or one sister cell could make PP and the other one a PN division (Figure 3K; lower panel). These results suggest that the clonal relationships of boundary cells influence their proliferative capacity but not their division mode.

**Figure 3.**
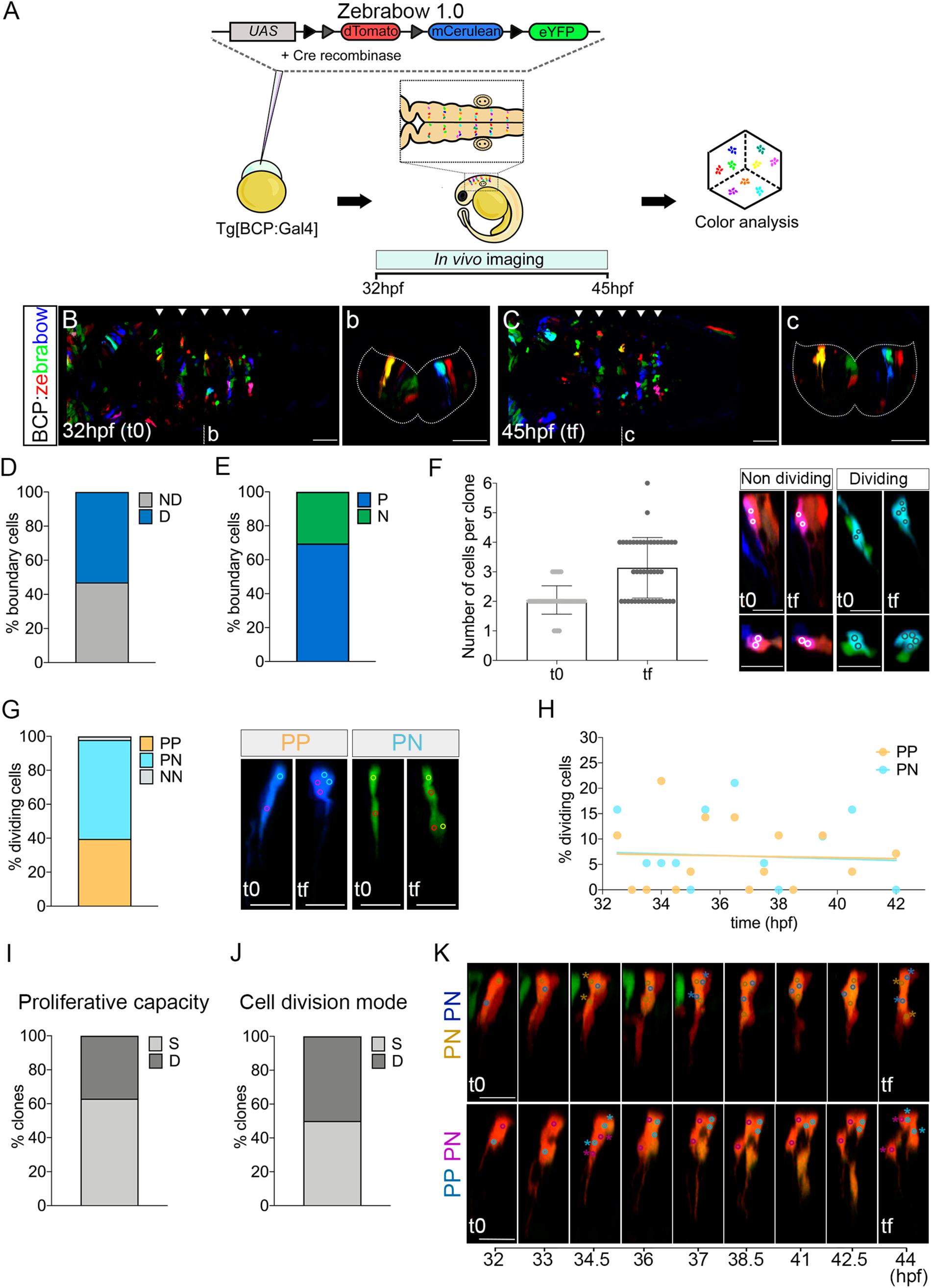
Multicolor clonal analysis of boundary cells. (A) Scheme depicting the experimental design to label and trace simultaneously the lineage of multiple boundary progenitors. Tg[BCP:Gal4] embryos at one cell stage were injected with Cre protein and the UAS:zebrabow-1.0 construct, which contains three fluorescent marker genes (dTomato, mCerulean, and eYFP). Hindbrains were *in vivo* imaged from 32 to 45hpf. (B–C, b–c) Dorsal MIPs and transverse projections through r4/r5 at t0 (32hpf) and tf (45hpf). White arrowheads indicate the position of the hindbrain boundaries. Dashed white line delimitates the contour of the neural tube. Scale bar 50μm. See Video EV1. (C) Proliferative capacity of boundary progenitors (53% dividing vs. 47% non-dividing, n=90). (D) Fate of boundary cell derivatives at 45hpf (tf). Out of 138 tracked boundary cells, 70% were progenitors (P), and 30% neurons (N). (E) Number of cells per clone at t0 and tf (t0, 2.1 ± 0.5 cells vs. tf, 3.1 ± 1 cells, n=44 clones, N=4 embryos). The plot shows the mean ± SD. Images show examples of non-dividing (white circled dots) and dividing (black circled dots) clones at 32hpf (t0) and 45hpf (tf). Top, transverse with ventricular surface to the top; bottom, dorsal view with anterior to the left. (F) Graph showing the cell division modes of boundary progenitor cells (n=48 cells): symmetric proliferative (PP, 40%), asymmetric (PN, 58%), and symmetric differentiative (NN, 2%). Transverse projections with ventricular surface to the top at t0 and tf showing examples of PP and PN divisions within boundary 2-cell clones. (G) Plot showing the percentage of boundary cells displaying PP (orange) or PN (turquoise) division mode over time (PP, 19 cells; PN, 28 cells; n=44 clones, N=4 embryos). Linear regression lines for PP (R square=0.002; p value=0.9 [ns]) and for PN (R square=0.004; p value= 0.8 [ns]) tend to zero. Note that PP and PN division modes are observed throughout the examined time window. (H–I) Clonal analysis of the proliferative capacity of sister cells and their cell division mode, respectively. Percentage of clones whose sisters display the same (S; 63%) or different (D; 37%) proliferative behavior (n=44 clones, N=4 embryos); and dividing boundary clones with sister cells displaying the same (S; 50%) or different (D; 50%) division mode (n=18 clones, N=4 embryos). (J) Examples of sister cells displaying the same (PN, PN) or different (PP, PN) division mode with corresponding time frames from t0 to tf. In the first panel, both the brown and the blue cells divide PN, the first at 34.5hpf and the second at 37hpf. In the second panel, both cells divide at 34.5hpf, the magenta cell undergoes a PN, the blue cell a PP division. Cell centers are circled and color-coded according to the lineage. BCP, Boundary Cell Population; hpf, hours post fertilization. Scale bar 20μm.

To assess whether Her9 controls the proliferative behavior of boundary cells, we performed functional multicolor clonal analysis (Loulier *et al*, 2014). We generated a new version of the zebrabow transgenes, zebrabow2.0, in which one of the color labels (red) coexpressed either the her9DN or the her9FL. This allowed us to specifically modulate Her9 in boundary cell clones identified by a specific color marker (in this case tdTomato), and compare their fate in the embryo to wild-type clones marked with distinct colors (Figure 4A; red clones expressing the her9DN or her9FL vs. non-red clones behaving as wild-type). In both loss- and gain-of-function experiments, we observed high color diversity (Figure EV5) and classified red and non-red clones in the same embryo at both 36 and 48hpf. When cell fate was addressed at 48hpf, her9DN clones showed a significant higher proportion of neurons while her9FL clones displayed more progenitors when compared to control clones (Figure 4B–C). These results reinforce the previous observation that Her9 promotes the maintenance of the pool of boundary progenitors.

**Figure 4.**
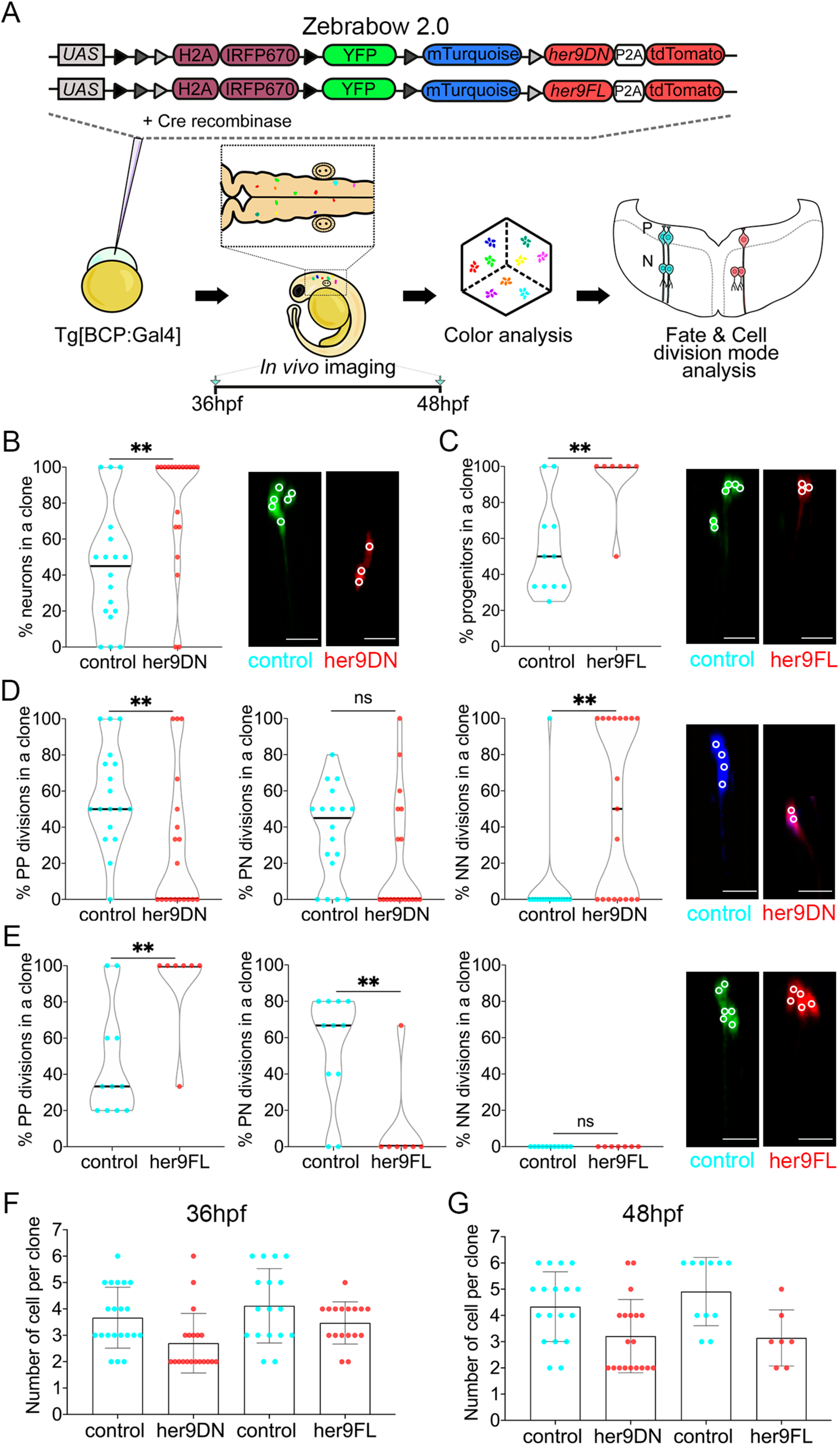
Her9 controls the fate and behavior of boundary cells. (A) Scheme depicting the experimental design. Tg[BCP:Gal4] embryos at one cell stage were injected with Cre protein and the UAS:her9DN-zebrabow2.0 or the UAS:her9FL-zebrabow2.0 constructs to either conditionally downregulate or activate Her9 in the boundary cells. Transgenes contain the YFP, mTurquoise, and tdTomato genes, and tdTomato-labeled clones coexpress either her9DN or her9FL in the context of the non-red wild-type clones. For color analysis, hindbrains were imaged *in vivo* at 36hpf and 48hpf. Then, the clones were analyzed for color, fate and division mode. (B–C) Boundary cell fate in *her9* loss-or gain-of-function at 48hpf. Violin plots show the percentage of neurons or progenitors in boundary control clones and upon her9DN or her9FL expression, respectively. Images show examples of control, her9DN, or her9FL boundary clones. (B) 43.4% ± 33 of neurons in control clones (median=45%) vs. 78.9% ± 33.7 in her9DN clones (median=100%), p value= 0.003 [**]; note that the her9DN clone displayed is only composed of neurons. (C) 53.8% ± 26.7 of progenitors in control clones (median=50%) vs. 92.9% ± 18.9 in her9FL clones (median=100%), p value=0.008 [**]; note that her9FL clone displayed is only composed by progenitors. (D) Boundary cell division mode in *her9* loss-of-function at 48hpf. Violin plots show the percentage of PP (left), PN (middle), and NN (right) divisions (PP: 57.4% ± 27.8 in control clones [median=50%] vs. 28.6% ± 37.7 in her9DN clones [median=0%], p value= 0.007 [**]; PN: 37% ± 25.5 in control clones [median=45%] vs. 21.4% ± 32.1 in her9DN clones [median=0%], p value= 0.06 [ns]; and NN: 5.6% ± 23.6 in control clones [median=0%] vs. 50% ± 47.5 in her9DN clones [median=50%], p value= 0.001 [**]). (E) Boundary cell division mode in *her9* gain-of-function at 48hpf. Violin plots show the percentage of PP (left), PN (middle), and NN (right) divisions (PP: 45.5% ± 30.7 in control clones [median=33.3%] vs. 90.5% ± 25.2 in her9FL clones [median=100%], p value=0.009 [**]; PN: 54.6% ± 30.7 in control clones [median=66.7%] vs. 9.5% ± 25.2 in her9FL clones [median=0%], p value=0.009 [**]; NN: 0% in control clones [median=0%] vs. 0% in her9FL clones [median=0%], p value>0.9 [ns]). All violin plots display the median in black. (F–G) Analysis of clonal cell growth upon *her9* loss- and gain-of-function at 36 and 48hpf, respectively. Plots show the number of cells per clone in each condition (mean ± SD): LOF: 3.7 ± 1.2 at 36hpf and 4.3 ± 1.3 at 48hpf in control clones vs. 2.7 ± 1.1 at 36hpf and 3.2 ± 1.4 at 48hpf in her9DN clones. GOF: 4.1 ± 1.4 at 36hpf and 4.9 ± 1.3 at 48hpf in control clones vs. 3.5 ± 0.8 at 36hpf and 3.1 ± 1.1 at 48hpf in her9FL clones. N of the analysis: 36hpf: n=21 control clones, n=20 her9DN clones (12 embryos); 48hpf: n=18 control clones, n=19 her9DN clones (15 embryos). 36hpf: n=17 control clones, n=17 her9FL clones (13 embryos); 48hpf: n=11 control clones, n=7 her9FL clones (15 embryos). Mann-Whitney test was performed in all compared conditions. All images display transverse projections of boundary clones with the ventricular surface towards the top. White circles in the images indicate cell centers. BCP, Boundary Cell Population; hpf, hours post fertilization; PP, progenitor-progenitor; PN, progenitor-neuron; NN, neuron-neuron divisions. Scale bar 20µm.

Next, we studied whether Her9 acts by modifying the boundary progenitors’ division mode. Boundary her9DN clones at 48hpf showed a decrease in the percentage of PP divisions and an increase of NN divisions in contrast to control clones (Figure 4D). In the same vein, sustained expression of *her9* resulted in an increase of PP and a decrease in PN divisions compared to controls at 48hpf (Figure 4E). These observations suggest that Her9 expands the boundary progenitor pool by maintaining PP divisions and preventing boundary cells from undergoing PN or NN divisions, and thus neurogenesis. Accordingly, when we analyzed clonal growth, we observed that already at 36hpf the her9DN clones were smaller compared to the control ones (Figure 4F), and this difference was maintained at 48hpf (Figure 4G). However, the size of control and her9FL clones at 36hpf was similar (Figure 4F). Noteworthy, at 48hpf her9FL clones displayed fewer boundary cells than controls (Figure 4G), indicating that her9FL clones did not grow during this temporal window. Overall, these results suggest that Her9 is necessary but not sufficient for the clonal growth of boundary cells.

### Her9 promotes the proliferation of boundary cells through *cdkn1ca*

To examine whether Her9 controls the proliferative capacity of boundary cells, we assessed the impact of *her9* downregulation in boundary cell proliferation at two embryonic stages, when *her9* expression was enriched in boundaries (28hpf), and at the time this expression had decreased (36hpf). We observed a lower number of boundary cells at 28hpf upon *her9* downregulation (Figure 5A–C). To determine whether this phenotype was caused by a decrease in the boundary cells entering in S-phase, we measured EdU-incorporation. We detected a lower percentage of EdU-positive boundary cells in her9-MO compared to control embryos (Figure 5D–F). Similar results were observed at 36hpf, with a decrease in the total number of boundary cells (Figure 5G–I). However, we found no differences in the percentage of cells engaged in S-phase (EdU-positive) upon *her9* downregulation (Figure 5J–L). These data indicate that Her9 controls the proliferative capacity of boundary cells at early embryonic stages.

**Figure 5.**
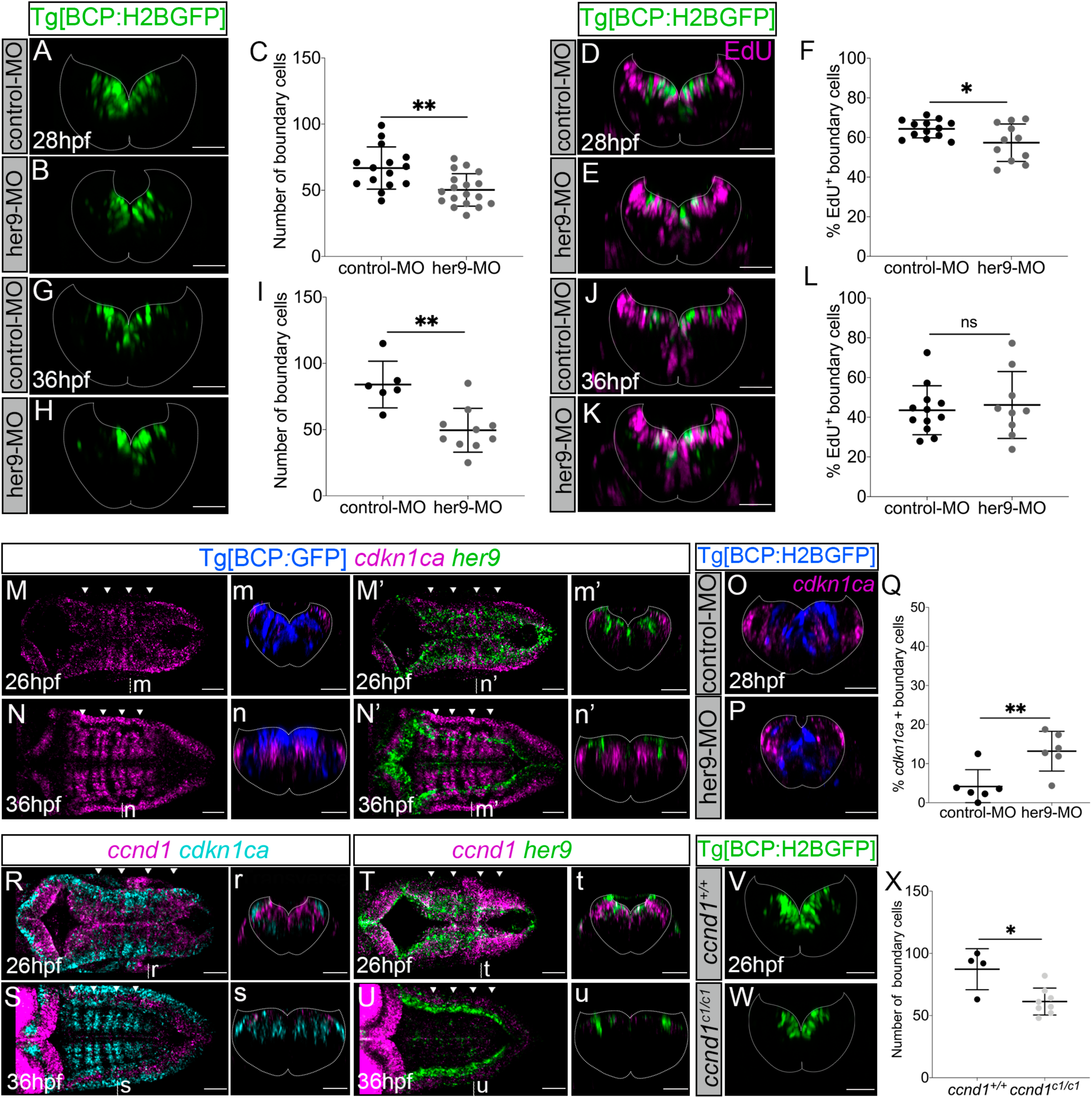
Her9 controls boundary cell proliferation through *cdkn1ca*. (A–B, D–E, G–H, J–K) Tg[BCP:H2BGFP] embryos injected with control-MO and her9-MO, displaying boundary nuclei in green at the indicated stages. (D–E, J–K) Embryos assayed for EdU-incorporation to detect S-phase cells (magenta). (C, I) Plots displaying the total number of boundary cells in r4/r5 at 28 and 36hpf, respectively (28hpf: control-MO: 66.8 ± 16 n=15 vs. her9-MO: 50.28 ± 12.3 cells n=18, p value=0.0029 [**], Welch’s test; 36hpf: control-MO 84 ± 17.6 n=6 vs. her9-MO 49.5 ± 16.5 n=10, p value=0.003[**], Welch’s test). (F, L) Plots showing the percentage of S-phase boundary cells at 28 and 36hpf, respectively (28hpf: control-MO 64.3% ± 4.5 n=13 vs. her9-MO 57.4% ± 9.5; n=11, p value=0.042 [*], Welch’s test; 36hpf: control-MO 43.5% ± 12.3 n=12 vs. her9-MO 46.17% ± 16.9 n=9; p value=0.695 [ns], Welch’s test). (M–N, M’–N’) *In situ* hybridization with *cdkn1ca* and *her9 in* Tg[BCP:GFP] embryos at indicated stages and displayed as single or double channels. (O–P) Tg[BCP:H2BGFP] embryos injected with control-MO or her9-MO and *in situ* hybridized for *cdkn1ca* at 28hpf. Boundary cells are displayed in blue. (Q) Plot displaying the percentage of boundary cells expressing *cdkn1ca* (4.1% ± 4.3 in control-MO n=6 vs. 13.2% ± 5.1 in her9-MO n=6; p value=0,009 [**], Mann-Whitney test). (R–S, T–U) *In situ* hybridization of *ccnd1* and with *cdkn1ca* or *her9* in Tg[BCP:GFP] embryos at the indicated stages. (V–W) Tg[BCP:H2BGFP;*ccnd1^+/+^*] and Tg[BCP:H2BGFP;*ccnd1^c1/c1^*] embryos at 26hpf with boundary cells in green. (X) Plot showing the number of boundary cells (87.3 ± 16.5 in *ccnd1^+/+^* n=4 vs. 61.3 ± 10.8 in *ccnd1^c1/c1^* n=8; p value=0.04 [*], Welch’s test). All images are transverse views of the r4/r5 boundary, except for (M–N, M’–N’, R–S, T–U) that are dorsal MIPs of the hindbrain with anterior to the left. The plots show the mean ± SD. Dashed white lines delimitate the contour of the neural tube. White arrowheads indicate the position of the hindbrain boundaries. BCP, Boundary Cell Population; hpf, hours post fertilization; MO, morpholino. Scale bar 50μm.

Hes/Her factors regulate the expression of cell cycle genes, and specifically of cell cycle arrest genes, in progenitor populations of different tissues (Georgia *et al*, 2006; Monahan *et al*, 2009; Zalc *et al*, 2014; Maeda *et al*, 2023). To determine whether Her9 controls boundary cell proliferation by the regulation of cell cycle genes, we first addressed the expression of *cdkn1ca*, a cell cycle arrest gene. *cdkn1ca* was expressed in the boundary flanking regions when engaged in neurogenesis, whereas it was absent in the boundaries when *her9* was highly expressed there (Figure 5M–N, M’–N’, m–n, m’–n’). Almost no boundary cells expressed *cdkn1ca* at early stages, whereas some started to express it after the loss of *her9* (Figure m–n, m’–n’). These results indicate that the onset of *cdkn1ca* expression follows the decline of *her9* in the boundaries and coincides with their commitment to neurogenesis. To seek whether *cdkn1ca* was the mediator of Her9 in the regulation of cell proliferation, we assessed *cdkn1ca* expression upon *her9* downregulation. We observed an increase in *cdkn1ca* expression in boundary cells compared to controls (Figure 5O–Q). Hence, when Her9 is highly expressed in boundary cells, it represses *cdkn1ca* expression in this cell population, promoting cell proliferation.

Cell cycle arrest proteins, such as Cdkn1ca and its orthologous p57, regulate cell proliferation through the inhibition of Cyclin/Cdk complexes (Grison & Atanasoski, 2020). Thus, we next studied the dynamics of the cell cycle progression CyclinD1 gene (*ccnd1*), which is expressed in hindbrain boundaries (Amoyel *et al*, 2005). *ccnd1* was enriched in hindbrain boundaries at early embryonic stages but greatly decreased by 36hpf in the whole hindbrain (Figure 5R–S, r–s). However, *ccnd1* did not overlap with *cdkn1ca* (Figure 5R–S, r–s). On the other hand, *ccnd1* and *her9* overlapped in boundary cells at 26hpf and their expression in the boundaries concomitantly decreased by 36hpf (Figure 5T–U, t–u). Therefore, *ccnd1* and *cdkn1ca* displayed complementary spatiotemporal patterns of expression, whereas *ccnd1* and *her9* overlapped in the hindbrain boundaries. To explore the putative role of CyclinD1 in the boundary cells proliferation, we generated a *ccnd1* loss-of-function mutant (Figure EV6A–B). Both *ccnd1^c1/+^* and *ccnd1^c1/c1^* embryos showed lower expression of the boundary-specific gene *sgca*, greatly in the most anterior hindbrain boundaries, compared to wild-type (Figure EV6C). To study whether this could indicate a partial loss of boundary cells, we analyzed the effect of *ccnd1* loss-of-function on boundary cell number. A significant reduction in the number of boundary cells was observed in *ccnd1^c1/c1^* mutants compared to *ccnd1^+/+^* embryos (Figure 5V– X). The decrease in the number of boundary cells was comparable to the one observed in *her9* morphants (compare Figure 5C and Figure 5X). Thus, CyclinD1 has an impact on the proliferative capacity of boundary cells. Overall, these data suggest that Her9 could control CyclinD1 activity in the boundaries through the downregulation of *cdkn1ca*, and thus, promote proliferation in boundary cells.

### *her9* is enriched in proliferating radial glial progenitors in a Notch independent manner at late embryonic stages

*her9* expression was temporary enriched in hindbrain boundaries. However later, *her9* expression was enriched in other territories of the hindbrain, including two lateral domains along the AP axis (Figures 1F and EV1C). To further explore what cell population expressed *her9* at later stages, we profiled it for the expression of *sox2* and *fabp7a,* markers of neural progenitors and radial glia, respectively. At 48hpf, *her9* expression colocalized with *sox2* along the ventricular domain, including the most medial and lateral domains, with few boundary cells still expressing it (Figure 6A–A’’, a–a’’; see asterisks). This scenario was maintained at 72hpf, where *her9* colocalized with *fabp7a* in the most medial and lateral domains (Figure 6B–B’, b–b’), indicating that these *her9*-cells were radial glial progenitors. We also detected comparable mitotic events between the *her9* positive lateral and medial progenitor population (6 ± 2.5 lateral vs. 8 ± 3.5, N=9 embryos; Figure 6B’; see white and gray arrows). Overall, *her9* is highly enriched in proliferating radial glia progenitors at the time neurogenesis ceases in the hindbrain. Next, we assessed whether *her9* expression in these domains depended on Notch-activity. When Notch-signaling was downregulated, we observed no differences in *her9* expression in these domains (Figure 6C–D), indicating that it did not rely on Notch activity. When we assessed the impact of *her9* downregulation, we detected a strong decrease in *sox2* expression in the whole hindbrain, including the lateral domains (Figure 6E–F; see asterisks). Taken together, our results suggest that Her9 plays a role in maintaining the stemness of distinct hindbrain progenitors’ populations in a Notch-independent manner at different temporal windows.

**Figure 6.**
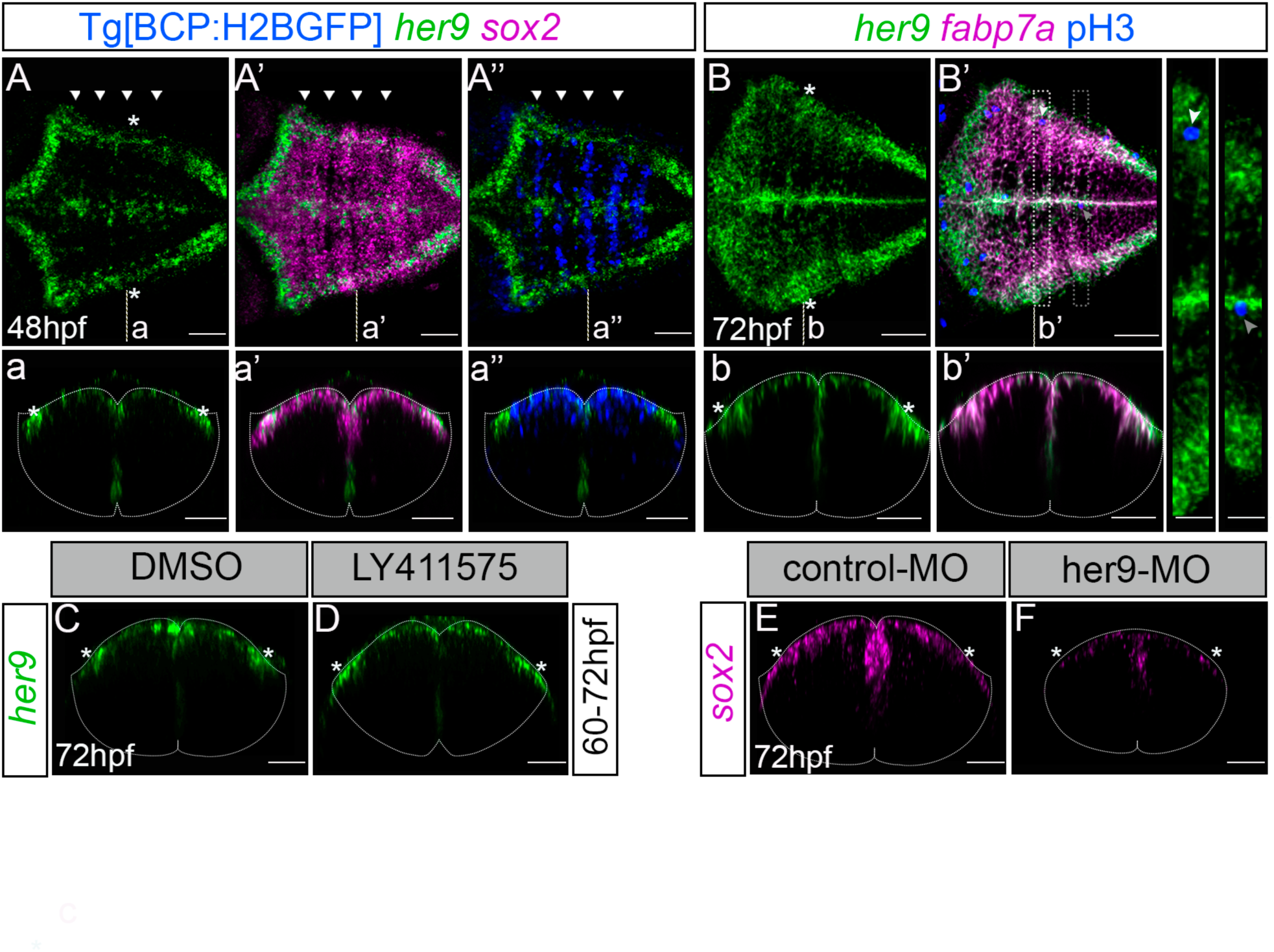
*her9* expression is maintained in different progenitor populations in the hindbrain at late embryonic stages. (A–A’’) Tg[BCP:H2BGFP] embryos at 48hpf *in situ* hybridized with *her9* and *sox2.* White arrowheads indicate the position of the hindbrain boundaries. (B–B’) Mü4127 embryos at 72hpf hybridized with *her9* and *fabp7a*, and immunostained with anti-pH3. Magnifications are indicated in the dorsal view with white and gray framed regions. White and gray arrows indicate the position of mitosis in lateral and medial domains, respectively. (A–A’’, B–B’) Dorsal MIP of the hindbrain with anterior to the left. Note that hindbrain lateral domains contain some proliferating radial glia progenitors (*fabp7a*) expressing *her9*. (C–D) Embryos treated with DMSO or LY411575 from 60hpf to 72hpf, and *in situ* hybridized with *her9* (DMSO n=7/7 vs. LY411575 n=6/6 embryos). Note that *her9* lateral domains of expression (asterisks) do not respond to Notch downregulation. (E–F) Embryos injected with control-MO and her9-MO, and *in situ* hybridized with *sox2* at 72hpf. Note *sox2* decreased in the her9-MO embryo (n=6/7) compared to the control (n=7/7). (a–a’’, b–b’, C–F) Transverse projections of r4/r5 boundary. White asterisks indicate the position of the *her9*-expressing lateral domains. BCP, Boundary Cell Population; hpf, hours post fertilization; MO, morpholino. Dashed white line delimitates the contour of the neural tube. Scale bar 50μm.

## DISCUSSION

Her9 maintains the stemness of boundary cells at the time these cells are specified, thus coinciding with the expression of the boundary genes (Figure 7A; (Cheng *et al*, 2004; Letelier *et al*, 2018). Her9 represses the neurogenic program by inhibiting *neurog1* and *neuroD4* (Figure 7B), consistent with the role of Her9 orthologue, Hes1, in the murine hindbrain (Baek *et al*, 2006). In absence of Notch activity in the hindbrain boundaries, Her9 expands the pool of progenitors by promoting symmetric proliferative cell divisions and preventing neurogenic divisions (Figure 7B). Later, *her9* loss coincides with the onset of Notch3-signaling and *her4* expression, promoting the transition of boundary cells to radial glia progenitors undergoing asymmetric divisions (Figure 7A, C). her9DN clones show an increase of symmetric neurogenic cell divisions at the expense of symmetric proliferative cell divisions as observed in compound *Hes* mice mutants (Hatakeyama *et al*, 2004). These observations fit into the model in which *Hes* genes maintain the stemness of neural progenitors before the onset of Notch activity (Hatakeyama & Kageyama, 2006). However, the moderate neuronal differentiation and increased Notch activity of boundary cells in the *her9* morphants suggest that Her9 could also prevent the onset of asymmetric cell divisions in hindbrain boundaries by repressing Notch activity through the inhibition of deltaD expression (Figure 7B–C). Our data in line with (Hevia *et al*, 2022) indicate that the boundary cell division mode is not determined by time, space, or clonal relationships, pointing towards a stochastic model of cell behavior after the onset of neurogenesis, similar to that of progenitor cells in the teleost retina (He *et al*, 2012). Nevertheless, the proliferative capacity of boundary cells seems to be deterministic, as clone size show little variability and sister cells tend to behave similarly in terms of cell proliferative capacity.

**Figure 7.**
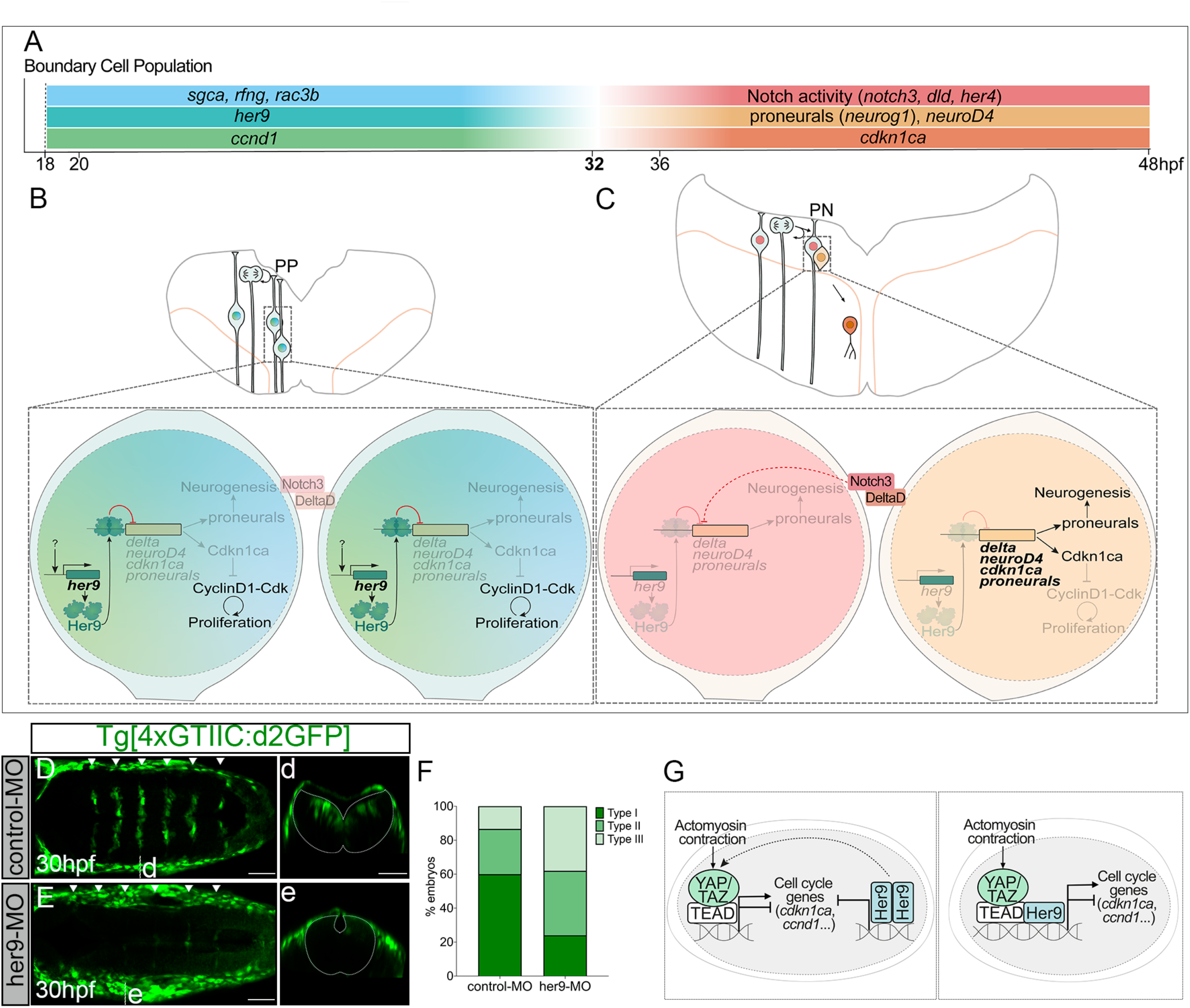
The strategy of stemness maintenance and expansion of hindbrain boundary cells. (A) Scheme of the temporal gene expression in the boundary cell population before and after 32hpf. (B–C) Scheme depicting transverse representations of the hindbrain boundaries before and after 32hpf. Magnifications (bottom) displaying the mechanisms of Her9 action. (B) Before 32hpf, Her9 inhibits neurogenic genes and cell cycle genes such as *cdkn1ca*, keeping boundary cells as progenitors dividing symmetric proliferative. (C) Upon *her9* decline, Notch3/DeltaD trigger asymmetric divisions. One cell is kept as a progenitor through Notch3 activity, whereas the other cell expresses both neurogenic genes and *cdkn1ca*, promoting neurogenesis and cell cycle exit. (D–E, d–e) Tg[4xGTIIC:d2GFP] embryos injected with control-MO or her9-MO displaying YAP/TAZ activity in green at 30hpf. Note that YAP/TAZ activity is inhibited upon *her9*-knowdown. (F) Graph with the percentage of embryos displaying the three phenotype strengths dependent in the effect of YAP/TAZ-activity in all boundaries (type I), less than 6 boundaries (type II), and in no boundaries (type III). 60% of embryos with type I (n=9/15), 26.7% type II (n=4/15), and 13.3% type III (n=2/15) in control-MO vs. 24% of embryos with type I (n=5/21), 38% type II (n=8/21), and 38% type III (n=8/21) in her9-MO). (G) Scheme of two scenarios of Yap/Taz-TEAD and Her9 cooperation in the control of boundary cell proliferation. In a first scenario, actomyosin contraction and Her9 may activate Yap/Taz-TEAD, and synergistically with Her9 it activates or inhibits the transcription of cell cycle related genes such as *cdkn1ca* and *ccnd1*. In a second scenario, actomyosin contraction activates Yap/Taz-TEAD, and Her9 binds to the Yap/Taz-TEAD complex activating or repressing the expression of cell cycle genes as *cdkn1ca* and *ccnd1*.

Her9 also promotes boundary cell proliferation by repressing *cdkn1ca* at early embryonic stages. In contrast, high levels of Hes1 promote low cell proliferation or quiescence (Baek *et al*, 2006; Shimojo *et al*, 2008; Sueda *et al*, 2019). Consistent with this, boundary cells are highly proliferative in teleosts as opposed to amniotes (Baek *et al*, 2006; Peretz *et al*, 2016; Voltes *et al*, 2019). This difference between high Her9 and high Hes1 in cell proliferation, could be explained by a newly described mechanism in which high Hes1 activates or inhibits proliferation of neural stem cells according to its shorter or longer expression time, resulting in the repression or activation of the cell cycle arrest protein p21, respectively (Maeda *et al*, 2023). *ccnd1* is also enriched in hindbrain boundaries and its mutation results in a lower number of boundary cells, suggesting that CyclinD1 controls their proliferative capacity (Figure 7A–C). Therefore, upon the inhibition of *cdkn1c*a by Her9, Cdkn1ca cannot repress CyclinD1 activity, promoting a faster cell cycle progression in boundaries compared to rhombomeric cells at early stages (Figure 7B). When Her9 decreases, boundary cells express Cdkn1ca that cannot block CyclinD1 activity and thus, they exit the cell cycle (Figure 7C). At early stages, Yap/Taz-TEAD activity expands boundary progenitors downstream of mechanical stimuli (Voltes et al., 2019). Moreover, our results showed that Yap/Taz-TEAD activity decreased in most hindbrain boundaries upon *her9* downregulation (30hpf; Figure 7D–F, d– e). One possible scenario is that Her9 could control Yap/Taz-TEAD activity in parallel to mechanical signals (Figure 7G). Alternatively, Yap/Taz-TEAD could form a transcriptional complex with Her9 downstream of mechanical signals as Hes1 does in the embryonic pancreas (Mamidi *et al*, 2018). In both cases, Her9 and Tap/Taz-TEAD would cooperate to promote cell cycle progression by repressing or activating cell cycle-related genes (Figure 7G; (Engel-Pizcueta & Pujades, 2021). Future functional studies should be performed to address their cooperation and regulation of the cell cycle in boundary cells.

*her9* expression is highly dynamic during hindbrain development. Its expression is enriched in boundary cells compared to rhombomere compartments at early stages, and it remains mainly in proliferating radial glial cells of lateral domains all along the AP axis at later stages. Our results support the developmental strategy othat different Hes/Her genes maintain distinct neural progenitor populations (Chapouton *et al*, 2006; Stigloher *et al*, 2008; Sigloch *et al*, 2023) establishing the neurogenic asynchrony in neural tissues. Hindbrain boundaries initially behave as a pool of non-neurogenic progenitors expressing *her9* in a Notch-independent manner. Later, hindbrain boundaries express *her4* and behave as a classical proneural cluster, as progenitors in telencephalic roof plate do (Dirian *et al*, 2014). Similar to boundaries, at later stages the remaining hindbrain progenitors in the lateral and medial domains express *her9* independently of Notch. These progenitors are greatly lost upon *her9* downregulation, and accordingly, Sox2 protein is enriched in the lateral domains when Notch activity ceases in the hindbrain (Hevia *et al*, 2022). Thus, we propose Her9 as a common player for maintaining distinct progenitor populations in the hindbrain at different temporal windows independently of Notch. Non-oscillatory expression of Hes1 correlates with high Hes1 and promotes stemness and low proliferation in neural progenitors (Baek *et al*, 2006; Shimojo *et al*, 2008; Sueda *et al*, 2019). Whether Her9 oscillatory dynamics could explain the differences between distinct hindbrain progenitor populations in space and time is an open question. Notably, derived mid-hindbrain boundary *her5* cells and telencephalic roof plate *her9/her6* progenitors show delayed neurogenesis and constitute a reservoir of neural stem cells in the adult brain (Chapouton *et al*, 2006; Dirian *et al*, 2014). Since there are very few boundary progenitors kept at later stages (Hevia *et al*, 2022) it seems unlikely that they would remain until adulthood. However, *her9*-positive domains of the hindbrain containing radial glial cells could harbor long-lasting progenitors until the adult stages. Future long-term lineage tracing of *her9* progenitors would be crucial to test this hypothesis.

## STRUCTURED METHODS

| Reagent/Resource | Reference or Source | Identifier / Catalog Number |
| --- | --- | --- |
| Experimental Models |  |  |
| Tg[elA:GFP] | (Labalette *et al*, 2011) | ZDB-TGCONSTRCT-110818-3 |
| Tg[HuC:GFP] | (Park *et al*, 2000) | ZDB-FISH-150901-19290 |
| Tg[BCP:Gal4] | (Hevia *et al*, 2022) | ZDB-ALT-230111-1 |
| Tg[UAS:GFP] | (Distel *et al*, 2009) | ZDB-ALT-100809-1 |
| Tg[cdh2:cdh2-tFT] | (Revenu *et al*, 2014) | ZDB-ALT-141218-4 |
| Tg[UAS:H2AmCherry] | (Hevia *et al*, 2022) | ZDB-ALT-200813-1 |
| Tg[tp1:d2GFP] | (Clark *et al*, 2012) | ZDB-ALT-120723-2 |
| Tg[UAS:H2BGFP] | (Hevia *et al*, 2022) |  |
| *ccnd1*<sup>c1/+</sup> | Generated in this work |  |
| Tg[4xGTIIC:d2GFP] | (Miesfeld & Link, 2014) | ZDB-ALT-150928-6 |
| Mü4127 | (Distel *et al*, 2009) | ZDB-ALT-120221-6 |
| Recombinant DNA |  |  |
| UAS:H2BCitrine | (Nikolaou *et al*, 2009) |  |
| H2BCitrine:UAS:her9DN | This study |  |
| H2BCitrine:UAS:her9FL | This study |  |
| UAS:zebrabow-1.0 | (Pan *et al*, 2013) |  |
| UAS:her9DN-zebrabow2.0<br>(in pUC57-mini vector) | GenScript Biotech (this study) |  |
| UAS:her9FL-zebrabow2.0<br>(in pUC57-mini vector) | This study |  |
| Antibodies |  |  |
| rabbit anti-GFP | Origene | TP401 |
| rabbit anti-phospho-Histone H3 (Ser10) | Upstate | 06-570 |
| anti-fluorescein-POD | Roche | 11426346910 |
| anti-digoxigenin-AP | Roche | 11093274910 |
| anti-digoxigenin-POD | Roche | 11207733910 |
| Donkey anti-Rabbit IgG (H+L) Highly Cross-Adsorbed Secondary Antibody, Alexa Fluor 488 | Invitrogen | A21206 |
| Goat anti-Rabbit IgG (H+L) Highly Cross-Adsorbed | Invitrogen | A21070 |
| Secondary Antibody, Alexa Fluor 633 |  |  |
| Oligonucleotides and other sequence-based reagents |  |  |
| PCR primers | This study | Table EV1 |
| 25-N random MO | Gene Tools, LLC |  |
| her9-2MO (referred to as her9-MO in the text) | (Bae *et al*, 2005) |  |
| p53-MO | (Langheinrich *et al*, 2002) |  |
| Chemicals, Enzymes, and other reagents |  |  |
| DMSO | CHEM-LAB | CL00.0422 |
| LY411575 | Sigma-Aldrich | SML0506-5MG |
| 1-phenyl-2-thiourea (PTU) | Sigma-Aldrich | P7629 |
| Tricaine | Sigma-Aldrich |  |
| DIG/FLUO RNA Labeling Mix | Roche | 11277073910/<br>11685619910 |
| RNA polymerase (SP6/T7) | Roche | 10810274001/<br>10881767001 |
| RRI (RNase inhibitor) | Takara | 2313 |
| Proteinase K | Sigma-Aldrich | P-2308 |
| Blocking Reagent | Roche | 14545800 |
| TSA solution (TSA Fluorescein/ TSA Cy3) | Perkin Elmer/Akoya | NEL753001KT |
| BM-Purple | Roche | 11442074001 |
| T3 mMessage mMachine Kit | Ambion Life Technologies | AM1348 |
| Golden Gate TALEN and TAL Effector Kit 2.0 | Addgene | Kit 1000000024 |
| SuperScript III kit | Invitrogen™ | 18080044 |
| TRIzol Reagent | Ambion Life Technologies | 15596018 |
| Qiagen® PCR Purification kit | Qiagen® | 28194 and 28106 |
| DreamTaq DNA polymerase | Thermo Scientific™ | EP0703 |
| Phusion™ High Fidelity DNA Polymerase | Thermo Fisher Scientific | F530S |
| XhoI | Thermo Scientific™ | FD0694 |
| NheI | Thermo Scientific™ | FD0974 |
| T4 DNA Ligase | Takara | 2011B |
| DH5α<sup>TM</sup> competent *E coli* | New England Biolabs | C2987H |
| BshTI (AgeI) | Thermo Scientific<sup>TM</sup> | ER1461 |
| XmaJ (AvrII) | Thermo Scientific<sup>TM</sup> | FD1564 |
| T4 DNA Ligase | New England Biolabs | M0202S |
| Bful | Thermo Scientific<sup>TM</sup> | ER1501 |
| Big Dye Terminator v3.1 | Applied Biosystems<sup>TM</sup> | 4337455 |
| Low melting agarose | Ecogen | AG0400 |
| Cre recombinase<br>(15,000 units/ml) | New England Biolabs | M0298M |
| Click-It<sup>TM</sup> EdU Alexa<br>Fluor<sup>TM</sup> 647 Imaging Kit | Thermo Scientific<sup>TM</sup> | C10340 |
| Software |  |  |
| GraphPad Prism 8 | GraphPad Software, Inc. |  |
| ImageJ | v2.3.0/1.53f51/FIJI |  |
| MaMut v.0.30.0 | (Wolff *et al*, 2018) |  |
| Photoshop | Adobe, version 23.5 |  |
| Illustrator | Adobe, version 27.3 |  |
| MatLab |  |  |
| Other |  |  |
| Glass-bottom Petri dishes | MatTek | P35G-1.5-14-C |
| FemtoJet Injector 4i | Eppendorf | 5252000021 |
| SP8 confocal | Leica |  |
| Macro Zoom Fluorescence<br>Microscope System<br>MVX10 | Olympus |  |

## METHODS AND PROTOCOLS

### Ethics declarations and approval for animal experiments

All procedures were approved by the institutional animal care, the PRBB Ethics Committee in Animal Experimentation and the Departament de Territori i Sostenibilitat (Generalitat of Catalonia) in compliance with the National and European regulations. Government and University veterinary inspectors audit the animal facilities and procedures to ensure that animal care standards are properly followed. The PRBB animal facility has the AAALAC International approval B9900073. All the members accessing the animal house must hold the international FELASA accreditation. The Project License covering the proposed work (Ref 10642, GC) pays particular attention to the 3Rs.

### Zebrafish strains

All zebrafish strains were maintained alternating generations of incrosses and outcrosses with wild-type. Embryos were obtained by mating adult fish following standard methods and grown at 28.5 or 23°C. 1% 1-phenyl-2-thiourea (PTU) (Sigma-Aldrich) was used as an inhibitor of pigmentation from 24hpf onward. Tg[BCP:Gal4], Tg[BCP:H2AmCherry], Tg[BCP:GFP], and Tg[BCP:H2BGFP] lines were used to label hindbrain boundary cells (Hevia *et al*, 2022). Tg[elA:GFP] and Mü4127 lines were used as landmarks of r3 and r5 (Labalette *et al*, 2011; Distel *et al*, 2009). Tg[HuC:GFP] line labels differentiated neurons (Park *et al*, 2000). The Tg[tp1:d2GFP] and Tg[4xGTIIC:d2GFP] lines monitor Notch-activity (Hevia *et al*, 2022; Park *et al*, 2000) and Yap/Taz-TEAD activity (Miesfeld & Link, 2014; Voltes *et al*, 2019), respectively. Tg[cadh2:GFP] line corresponds to the tandem fluorescent cadherin timer, which allows to *in vivo* monitor cadherin2 subcellular location (Revenu *et al*, 2014).

### TALEN-genome editing

The *ccnd1^c1^* line was generated using TALEN-induced mutagenesis strategy. A target site in the second exon and the corresponding left and right TALENs were designed using the online software MoJo Hand (http://www.talendesign.org). The TALEN repeat arrays were generated following the protocol described in (Cermak *et al*, 2011), and are available on the Addgene website (https://www.addgene.org). The array plasmids were fused to Fok1 endonuclease and linearized. mRNAs were *in vitro* transcribed using the T3 mMessage mMachine Kit (Ambion). The mRNAs of the left and right arms (1:1) were injected into one-cell stage embryos. gDNA of a subset of embryos from each clutch was extracted. The efficiency of the TALEN pair was assessed by amplifying a 550bp PCR fragment containing the target sites (Table EV1) and then digesting with BfuI that cut in the spacer of the target site (Figure EV6). Mosaic fish were outcrossed and a pool of embryos was genotyped as described to identify potential mutations. F1 fish were genotyped using fin clips and the PCR band carrying the mutation was sequenced. The *ccnd1^c1^* mutation consisted of a deletion of 2 nucleotides at position c.35 to c.36 replaced by T (Figure EV6A). This change causes a premature STOP codon generating a truncated CyclinD1 protein of 12AA without the cyclin box domain. To study the effects of the mutation in the boundary cells, *ccnd1^c1/+^* were outcrossed with Tg[BCP:H2BGFP] fish, and the progeny was raised to adulthood. Sibling embryos obtained from Tg[BCP:H2BGFP;*ccnd1^c1/+^*] outcrossed with *ccnd1^c1/+^* fish were *in vivo* imaged and genotyped as previously described (Figure EV6B).

### Pharmacological treatments

Embryos were dechorionated and treated with either 10μM of the gamma-secretase inhibitor LY411575 (Sigma-Aldrich) as an inhibitor of Notch-signaling or DMSO as a control, diluted in embryo medium. Embryos were incubated during the indicated temporal windows at 28.5°C. After treatment, embryos were washed with embryo media and fixed in 4%PFA for 3h at RT or O/N at 4°C.

### EdU-incorporation experiments

Cells in the S-phase were detected by EdU incorporation using the Click-It^TM^ EdU Alexa Fluor^TM^ 647 Imaging Kit (C10340, Thermo Fisher Scientific) according to (Belmonte-Mateos *et al*, 2023). Briefly, Tg[BCP:H2BGFP] embryos were dechorionated, incubated in 500µM EdU diluted in 7% DMSO fish water, and placed shaking on ice for one hour for better EdU incorporation. They were washed with embryo medium and fixed in 4%PFA O/N at 4°C. Embryos were permeabilized with proteinase K (10 mg/ml, Invitrogen), post-fixed, and washed in PBT. Then, they were incubated for 1h in 1%DMSO/1%Triton X-100/PBS. The Click-iT reaction was carried out according to the manufacturer’s instructions before immunostaining.

### Whole mount *in situ* hybridization

Embryo whole mount *in situ* hybridization was adapted from (Thisse & Thisse, 2008). The following antisense riboprobes were generated by *in vitro* transcription from cloned cDNAs: *deltaD* (Haddon *et al*, 1998)*, her9* (Leve *et al*, 2001), *neurod4* (Park *et al*, 2003), *neurog1* (Itoh & Chitnis, 2001), *sox2* (März *et al*, 2010), and *sgca* and *rfng* (Letelier *et al*, 2018). The other antisense probes were generated by PCR amplification adding the promoter sequence T7 or Sp6 in the reverse primers (Table EV1). Embryos were dehydrated and posteriorly rehydrated before permeabilization with proteinase K (10mg/ml, Invitrogen) within a range of 5 to 30 min according to the stage (18 to 72hpf). They were incubated in FLUO-[1:50] and DIG-labeled probes [1:100] diluted in hybridization buffer. Embryos were then incubated with an anti-FLUO-POD ([1:400], Roche) or anti-DIG-AP ([1:1000], Roche) in 2% blocking reagent (Roche), 10% Neutralized goat serum in 1x malic acid buffer in PBT (MABT) blocking solution, followed by anti-DIG-POD ([1:400], Roche). FLUO-and DIG-labelled probes were stained with TSA Fluorescein and Cy3 (Akoya), respectively. For the chromogenic *in situ* hybridization, DIG-labelled probes were stained with BM-Purple (Roche).

### *In toto* embryo immunostainings

Embryos were permeabilized with proteinase K (10mg/ml, Invitrogen) in 1% PBS, 0,1% Tween20 (PBT), post-fixed with 4%PFA, and blocked in 10% neutralized goat serum and 2% Bovine Serum Albumin (BSA) in PBT for 2h at RT. For immunostainings after *in situ* hybridization, embryos were blocked in 5% neutralized goat serum in PBT for 1h and incubated O/N at 4°C with the corresponding primary antibody: rabbit anti-GFP ([1:400], Torrey Pines), rabbit anti-pH3 ([1:200]; Upstate). After washings with PBT, embryos were incubated with secondary antibodies conjugated with Alexa Fluor^®^488 or 633 ([1:500], Invitrogen).

### Confocal imaging of whole mount embryos

Embryos were mounted in either 1% low melting agarose (Ecogen) or 0.7% for time-lapse imaging, with the hindbrain towards the glass of the glass-bottom Petri dishes. Low-melting agarose was dissolved in embryo water with 1% PTU (Sigma-Aldrich) or PBT for live and fixed embryos, respectively. Live embryos were anesthetized with 0.1% tricaine (Sigma-Aldrich). All images were acquired in a SP8 Leica inverted confocal microscope. Different combinations of 458, 488, 514, 561, and 633nm lasers were used to excite fluorochromes and emitted light was detected with PMT or HyD detectors. Each channel was acquired by line in live embryos by stacks in fixed embryos. Images of live embryos were acquired with a 20x immersion objective with glycerol oil (NA 0.7, z-step 1.19µm). The 20x dry objective (NA 0.75, z-step 0.79 µm) was used for all fixed samples except for her9-MO experiments that were acquired with 20x immersion objective with glycerol oil (NA 0.75). The image format was 1024x512 and the scan speed ranged from 400 to 600Hz. The software zoom was between 0.85x–1.65x.

### Morpholino knockdown experiments

Embryos were injected at the one-cell stage with: i) 8ng random 25N morpholino as control (Gene Tools, LLC); or ii) 8ng splicing-blocking morpholino her9-2MO (her9-MO in the text; (Bae *et al*, 2005). 7.5ng p53-MO (Langheinrich *et al*, 2002) was included in all MO-injections to diminish putative artifacts (Gerety & Wilkinson, 2011). Embryos were grown at 28.5°C until the desired stage. To assess her9-MO efficiency, we analyzed the fourth ventricle opening defects (Bae *et al*, 2005). The penetrance of this phenotype was around 80% (control-MO N=0/8 vs. her9-MO N=10/13 embryos; Figure EV2B–C, b–c), although some degree of variability was observed between independent experiments. We checked as well for the presence of the *her9* spliced-defective variants in the same control-MO and her9-MO embryos at 36hpf (Figure EV2D). RNA of the two pools of embryos was extracted with TRIzol reagent (Ambion) and phenol/chloroform protocol. The RT-PCR was performed with the SuperScript III kit (Invitrogen) following the manufacturer’s instructions. The MO binding site was amplified with the primers in Table EV1.

### Single clonal color analysis

Constructs for *her9* loss- and gain-of-function analyses contained the UAS sequence (H2BCitrine:UAS:her9DN/her9FL) to conditionally alter Her9’s function specifically in hindbrain boundary cells using the Tg[BCP:Gal4] (Hevia *et al*, 2022). For loss-of-function experiments, a dominant negative form of her9 was generated by removing the WRPW domain, necessary for Her9’s transcriptional repressor activity (her9^ΔWRPW^, her9DN in the text; (Fisher *et al*, 1996; McLarren *et al*, 2001). For gain-of-function, we cloned the full-length version of the *her9* gene (her9FL). For single-color analyses, her9FL and her9DN were amplified by PCR from *her9* cDNA plasmid (Leve *et al*, 2001) with Phusion polymerase (Thermo Scientific) using the primers in Table EV1. PCR products and the H2BCitrine:UAS plasmid (Nikolaou *et al*, 2009) were digested with XhoI and NheI, and ligated with T4 DNA ligase (Takara) in a 5:1 (insert:vector) ratio. Next, the H2BCitrine:UAS, H2BCitrine:UAS:her9FL or H2BCitrine:UAS:her9DN were microinjected in Tg[BCP:Gal4] embryos at one-cell stage at 15ng/μl with Tol2 mRNA at 18ng/μl in a 1nl drop. For the analysis, the number of cells expressing H2BCitrine in each boundary were annotated. In H2BCitrine:UAS:her9DN/her9FL conditions, we quantified the percentage of boundary cells expressing each gene of interest over the number of H2BCitrine-cells exhibiting overexpression or ectopic expression of *her9*.

### Multicolor clonal tracking and analysis

#### Design and cloning

For functional multicolor clonal analysis, we designed two zebrabow2.0 UAS constructs co-expressing either her9DN or her9FL with one of the fluorescent proteins of a quadrichromatic brainbow cassete as modified from that described in (Loulier *et al*, 2014). The two zebrabow2.0 constructs (UAS:her9DN-zebrabow2.0 and UAS:her9FL-zebrabow 2.0) express by default the far-red protein iRF670 (Shcherbakova & Verkhusha, 2013) fused to the zebrafish histone gene *h2az2a* (Wilkinson & Shyu, 2001) to direct expression in the nucleus; upon Cre recombinase action, expression switches to one of the three spectrally distinct fluorescent proteins: mEYFP (Zacharias *et al*, 2002), mTurquoise2 (Goedhart *et al*, 2012), or tdTomato (Shaner *et al*, 2004). her9DN and her9FL were separated from the tdTomato by the zebrafish cleavage peptide P2A (Kim *et al*, 2011). These transgenes enable equilibrated expression of 3 recombination outcome independently of Cre dosage and allow the comparison of the behavior of her9DN-or her9FL-expressing clones (tdTomato-positive) with wild-type clones (tdTomato-negative) within the same embryo. The designed sequence was synthesized in a pUC57-mini vector (GenScript Biotech). This plasmid was also designed as a versatile functional tool to allow the insertion of any gene of interest and the modification of the different fluorescent proteins by restriction enzymes. For the *her9* gain-of-function (UAS:her9FL-zebrabow2.0), her9FL was amplified as described above using primers listed in Table EV1, and cloned within BshTI and XmaJ sites.

#### Injection

Tg[BCP:Gal4] embryos at one-cell stage were microinjected with UAS:zebrabow-1.0 (Pan *et al*, 2013), UAS:her9DN-zebrabow2.0 or UAS:her9FL-zebrabow2.0 (see before) at 10ng/μl with Tol2 mRNA at 18ng/μl and 0.25μl Cre recombinase at 15.000 units/ml (NEB) in 4nl drop as (Brockway *et al*, 2019). For multicolor functional experiments, we injected 0,5μl of Cre to increase recombination efficiency.

#### Imaging

Embryos were selected and imaged *in vivo*. For the wild-type clonal study, live embryos were imaged every hour from 32 (t0) to 48hpf (tf) at 23°C in independent imaging sessions. The embryonic stage was corrected according to the temperature used for imaging. For the functional study, embryos were *in vivo* imaged at 36 and 48hpf. Between imaging sessions, embryos were demounted and grown at 23°C. In these experiments, we also acquired the bright field channel to observe the contour and landmarks of the hindbrain.

#### Image processing

Images were rotated to align the hindbrain midline with the X-axis. In time-lapses, a drift correction was performed using a modified version of the script used in (Dyballa *et al*, 2017). A variable ROI was used to include the whole hindbrain boundary in all time frames (time-lapses) or individual clones at fixed points (functional analysis). Images were then converted to .xml/.hdf5 using the Big Data Viewer Plugin.

#### Color analysis

To establish the clonal criteria in the multicolor time-lapse data, we analyzed color frequency to identify clones with less frequent colors to define the characteristics for all tracked clones. Cell centers within boundary ROIs were pointed and saved using ROI Manager in ImageJ. An average of red, green, and blue channel values was determined through semi-manual ROI measurements and a ternary plot was generated using ImageJ macros adapted from (Loulier *et al*, 2014). In all embryos, the number of cells in each of the 25 color identities defined according to the subdivisions of the RGB space was quantified (Figure EV4A). Color identities higher than 10% were considered frequent, whereas colors lower than 2.5% were considered rare (Figure EV4B–C). Based on the size and fate of rare color clones, we followed boundary cell clones located in the progenitor domain at t0 and containing one to three cells per clone. Color analysis was further used to confirm the clonal relationship of cells. Cells falling in the same RGB subdivision and in close contact were considered clonally related (Figure EV4A–B). Boundary clones with frequent colors were only followed when isolated. To represent the applied color and the spatial criteria followed, we generated a 3D-ternary plot in the ML axis using a Matlab code (Figure EV4A; (Loulier *et al*, 2014). For the functional study, we followed the same color analysis to establish clonal relationship, and the mean intensity of the red channel to classify her9DN/her9FL (red) and control (non-red) clones. Clones were considered red when displaying more than 5% of red intensity, and non-red when displaying less than 1% (Figure EV5A–B). Clones that did not fall into these categories were excluded from the analysis.

#### Cell counting and tracking

For the time-lapses, cells were annotated and tracked in MaMut software v.0.30.0 (Wolff *et al*, 2018) from t0 to tf. For these analyses, we tracked 44 clones in different hindbrain boundaries of four different embryos (r2/r3, r3/r4, r4/r5 and r6/r7 in e10_2205; r2/r3, r3/r4, and r4/r5 in e1_3005; r3/r4, r4/r5, and r5/r6 in e3_1505; and r4/r5 in e7_2503). A number was ascribed to each clone in a boundary according to its ML position (from left to right). All clones, cell lineages, and cell divisions were displayed as lineage trees obtained from the MaMut Track Scheme and posteriorly drawn in Illustrator (Adobe) (Figure EV4D). For the functional analyses, cells were annotated in MaMut software v7.0.2 (Wolff *et al*, 2018). We analyzed a total of 12 embryos at 36hpf (21 non-red vs. 20 red clones) and 15 embryos at 48hpf (18 non-red vs. 19 red clones) of *her9* loss-of-function condition; and 13 embryos at 36hpf (17 non-red vs. 19 red clones) and 11 embryos at 48hpf (11 non-red vs. 7 red clones) of *her9* gain-of-function. Clones were ascribed a number in each boundary according to their ML and dorsoventral (DV) position. The same number was used to link the data of the same clone from 36 to 48hpf.

#### Cell fate, proliferative capacity, and cell division modes analyses

For clonal analyses we annotated: i) the number of cells at t0 and tf; ii) the number and time of cell divisions per clone in the wild-type tracked clones; iii) the cell fate at tf (progenitor, P vs. neuron, N) by the relative position to the ventricle and by the presence of the apical contact (a cell was considered a progenitor when the nucleus was close to the ventricle and showed an apical contact, or a neuron if the nucleus was close to the mantle zone and had no apical contact; Figure EV4A); and iv) the cell division mode (PP, PN, NN). The cell division mode was determined according to the defined fate and relative position of daughter cells. A division was considered PN when the two daughter cells were not in close contact, with one closer to the ventricle and the other to the mantle zone; and a division was considered PP or NN when both daughter cells were in close contact and located at the same DV level. For the cell division mode analysis across time, fate was not ascribed for cells that were tracked less than 3h after division. In the functional analysis, the same criterion was followed. The number of PP divisions in a clone was defined by the number of progenitors in the clone, as every PP division provides one new progenitor. The number of PN in a clone was established by the number of neurons in the clone, as every PN division provides one new neuron. Clones only composed of an even number of neurons at 36hpf were considered to undergo NN divisions.

### Quantification and statistical analyses

#### Cell counting

We defined the independent experimental units according to the experimental set ups: i) embryos in knockdown (morpholino) experiments, in which we analyzed the effects at the whole cell population level; ii) hindbrain boundaries in the single-color analysis, in which we cannot segregate single clones; and iii) clones in the multicolor functional analysis, in which we examined single and isolated clones. We quantified the number of boundary cells in the r4/r5 boundary in all imaged embryos, except for single-color and multicolor clonal analyses in which we analyzed r2/r3, r3/r4, r4/r5, and r5/r6 boundaries. Cells were counted within a ROI defined for each experiment (the same in control and experimental conditions) and established by the mean of the ROIs including a whole boundary of all the embryos. We counted the cells using the MaMut FIJI Plugin v7.0.2 (Wolff *et al*, 2018) and annotated the number of boundary cells expressing the gene or marker of interest.

#### Statistical analyses

We performed either normality tests and the corresponding t-test (Welch’s test) or non-parametric test (Mann-Whitney test) according to the distribution of the data. Values were expressed as mean ± SD. All graphs were generated with GraphPad Prism 8 software. Image brightness and contrast were linearly adjusted in ImageJ. Median filters were applied in images in Figures 1 and Figure EV1 only for presentation purposes. Bleach Correction Plugin of ImageJ was used for the red channel in the multicolor time-lapse example (Video 1) for better representation. All figures were assembled in Photoshop (Adobe) and illustrations were done in Illustrator (Adobe).

## ACKNOWLEDGEMENTS

We would like to thank Laia Subirana, Clàudia Garcia, and Dulce Real for technical assistance and members of the lab for insights and critical discussions. We thank TA Weissman for the UAS:zebrabow1.0 plasmid, and V Lecaudey for her help in the generation of *ccnd1* mutants and her critical insights on the manuscript. We also want to thank the members of Jean Livet’s lab for their help. Confocal microscopy was performed at the Advanced Light Microscopy Unit at the CRG. This work was funded by grants PGC2018-095663-B-I00 and PID2021-123261NB-I00 from Ministerio de Ciencia y Universidades (MICIU), Agencia Estatal de Investigación (AEI, DOI: 10.13039/501100011033) and Fondo Europeo de Desarrollo Regional (FEDER) to CP. The Department of Medicine and Life Sciences (UPF) is a Unidad de Excelencia María de Maeztu (CEX2018-000792-M) funded by the AEI. This work was also supported by IHU FOReSIGHT ANR-18-IAHU-01 to JL. CEP was a recipient of FPU predoctoral fellowship from the MICIU and was awarded with the short Ph.D. stay fellowship of FPU from the MICIU. AV was a recipient of a predoctoral fellowship from LaCaixa Foundation. CP is a recipient of ICREA Academia award (Generalitat de Catalunya).

## Disclosure and competing interest’s statement

The authors declare no conflict of interests.

## AUTHOR’S CONTRIBUTIONS

Conceptualization: CEP, CFH, and CP; Methodology: CEP, CFH, and CP; Investigation and formal analysis: CEP; Validation: CEP, CFH, JL, and CP; Resources: AV, and JL; Design and analysis of the multicolor clonal studies: CEP and JL; Writing-original draft: CEP and CP; Writing-review & editing: CEP, CFH, JL, and CP; Visualization: CEP; Supervision: CP; Project administration: CP; Funding acquisition: CP.

## EXPANDED VIEW FIGURE LEGENDS

**Figure EV1.**
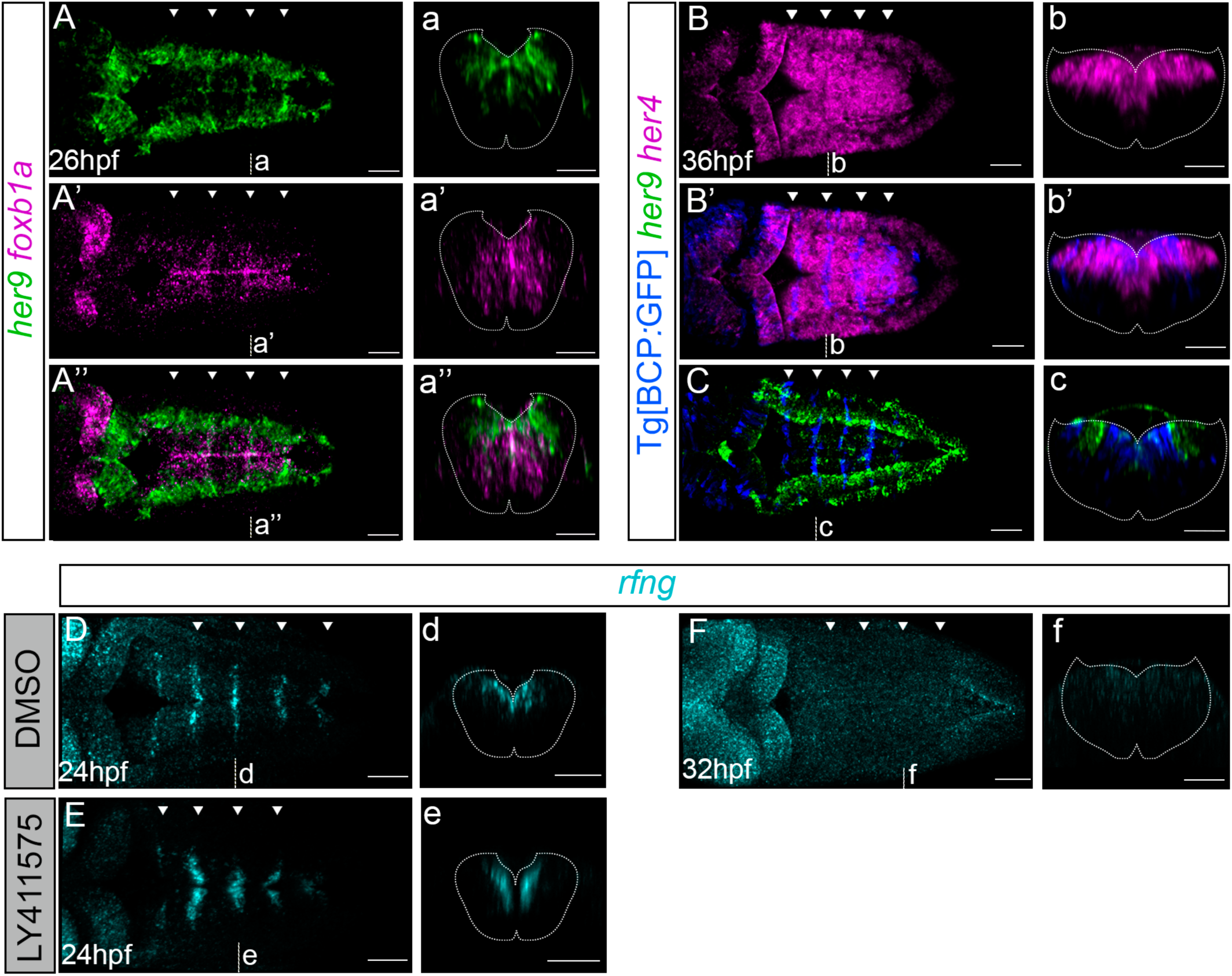
Enriched *her9* expression overlaps with hindbrain boundary markers. (A–A’’) *In situ* hybridization of *her9* and the *foxb1a* boundary marker in Tg[BCP:GFP] embryos at 26hpf. Dorsal views of single (A–A’) or merged channels (A’’). (a–a’’) Transverse projections of (A–A’’) through r4/r5 boundary. (B–B’, C) *In situ* hybridization of *her4* or *her9* in Tg[BCP:GFP] embryos at 36hpf. Dorsal views of single (B) or merged channels are shown (B’, C). (b–b’, c) Transverse projections of (B–B’, C) through r3/r4 boundary. (D–E) Embryos treated from 18hpf to 24hpf either with DMSO (D) or with LY411575 (E) and *in situ* hybridized with *rfng*. (d–f) Transverse five stack projection (D–F) through r4/r5. (F) *In situ* hybridization of *rfng* in Tg[elA:GFP] embryos at 32hpf. (f) Transverse five stack projection (F) through r4/r5. White arrowheads indicate the position of the hindbrain boundaries. Dashed white line delimitates the contour of the neural tube. BCP, Boundary Cell Population; hpf, hours post fertilization. Scale bar 50μm.

**Figure EV2.**
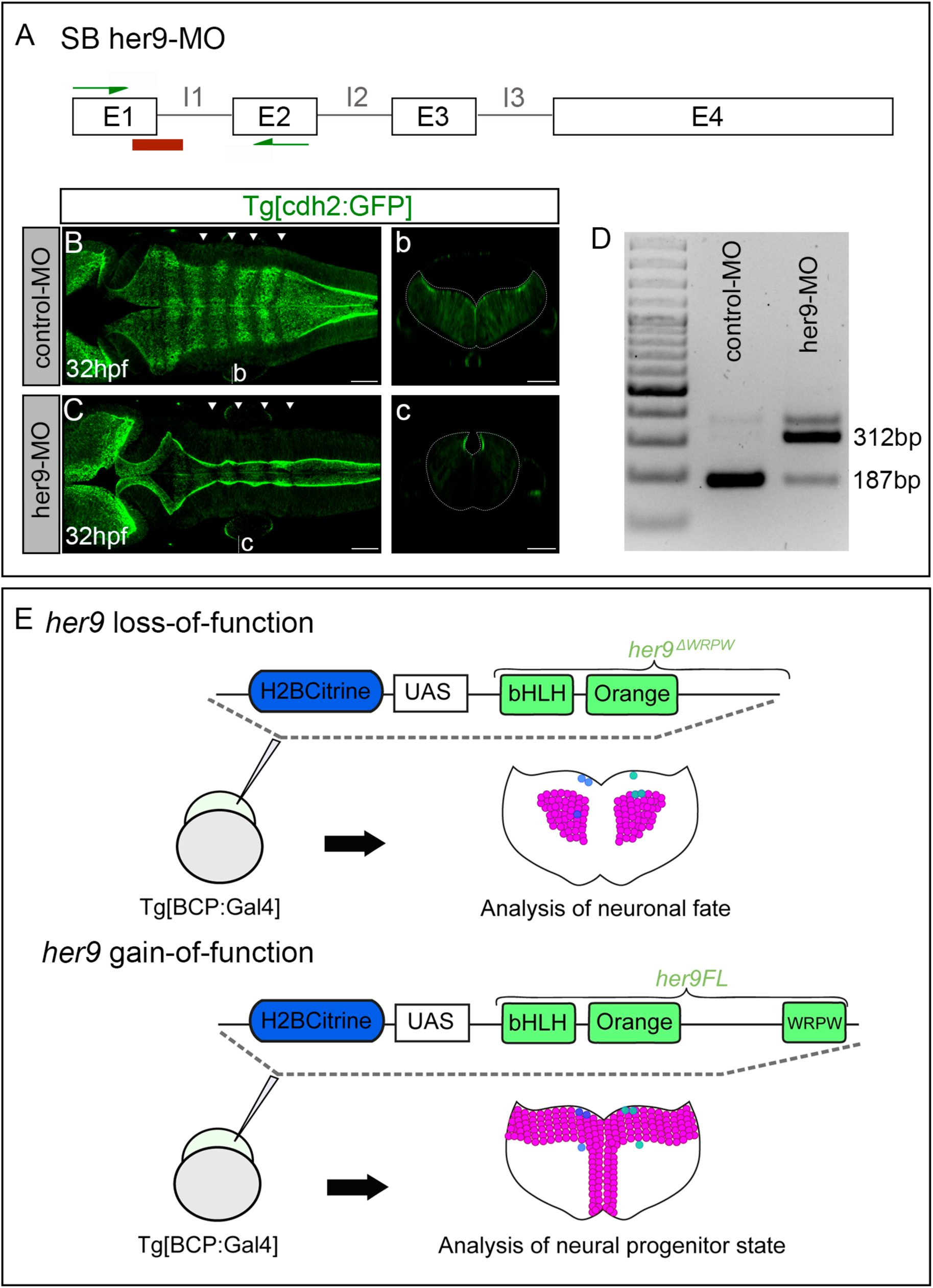
Loss-and gain-of-function of *her9*. (A) Scheme depicting the structure of the splicing blocking *her9* morpholino, SB her9-MO. Position of the her9-MO (red box), and the primers used to assess its efficiency (green arrows). E1–E4, exon 1–4; I1–I3, intron 1–3. (B–C, b–c) Tg[cdh2:GFP] embryos injected with either control-MO or her9-MO at one cell stage. Note that Her9 downregulation resulted in defects in the opening of the hindbrain ventricle (n=10/13), whereas no defects were observed in controls (n=8/8) at 32hpf. (B–C) Dorsal views with anterior to the left, (b–c) transverse five stack projections of (B–C) through r4/r5 boundary. White arrowheads indicate the position of the hindbrain boundaries. Dashed white line delimitates the contour of the neural tube. Scale bar 50μm. (D) Agarose gel showing the *her9* spliced-defective variant detected by RT-PCR in control-MO or her9-MO embryos in (B–C). (E) Scheme depicting the conditional loss- and gain-of-function experiments. Tg[BCP:Gal4] embryos at one cell stage were injected with either H2Bcitrine:UAS:her9DN (LOF, her9^ΔWRPW^) or H2Bcitrine:UAS:her9FL (GOF) to specifically express them in boundary cells. Embryos were let to develop until 36hpf when analysis of progenitor vs. neuronal cell fate was performed.

**Figure EV3.**
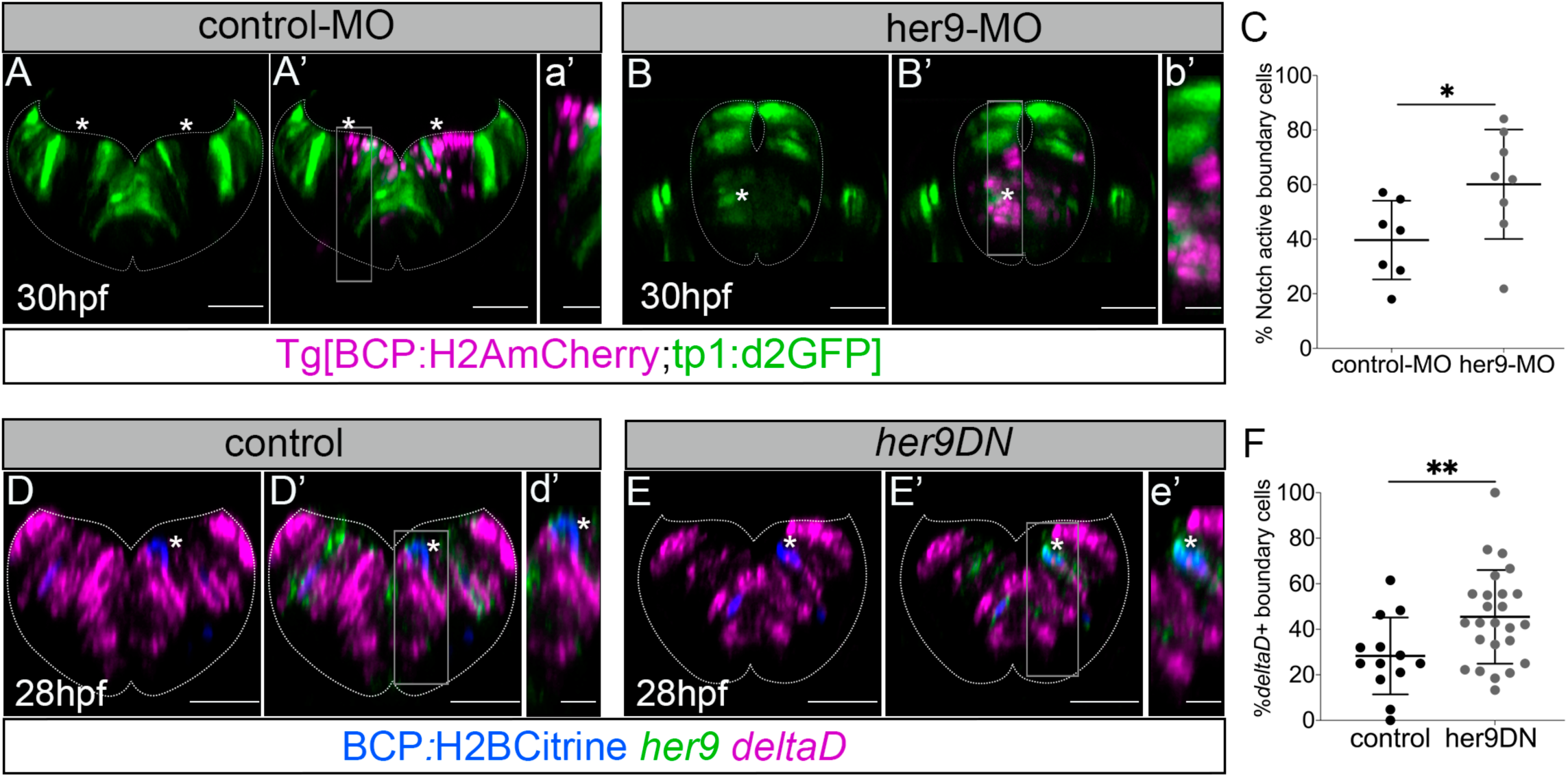
Her9 prevents the onset of Notch activity at early embryonic stages. (A–A’, B–B’) Tg[BCP:H2AmCherry;tp1:d2GFP] embryos injected with control-MO or her9-MO, displaying Notch activity in green and the boundary cell nuclei in magenta at 30hpf. Transverse five stack projection of r4/r5. (C) Plot showing the Notch active boundary cells in control-MO and her9-MO embryos (39.7% ± 14.4 in control-MO n=7 vs. 60.1% ± 20.1 in her9-MO n=8; p value=0.04 [*], Welch’s test). (D–D’, E–E’) Tg[BCP:Gal4] embryos injected with H2Bcitrine:UAS or H2Bcitrine:UAS:her9DN and *in situ* hybridized with *her9* and *deltaD*. Boundary cells (H2Bcitrine) are displayed in blue. (D–D’, E–E’) Transverse single-sections of r3/r4 or r4/r5, respectively. (a’–e’) Magnifications of the gray framed regions in (A’–E’). Dashed white line delimitates the contour of the neural tube. (F) Plot displaying the percentage of boundary cells expressing *deltaD* (28.3% ± 16.9 in control embryos, n=13 boundaries, N=6 embryos vs. 45.5% ± 20.6 in her9DN embryos, n=25 boundaries, N=10 embryos; p value=0.009 [**], Welch’s test). BCP, Boundary Cell Population; hpf, hours post fertilization; MO, morpholino. The plots show the mean ± SD. Scale bar 50μm and 20μm for magnifications.

**Figure EV4.**
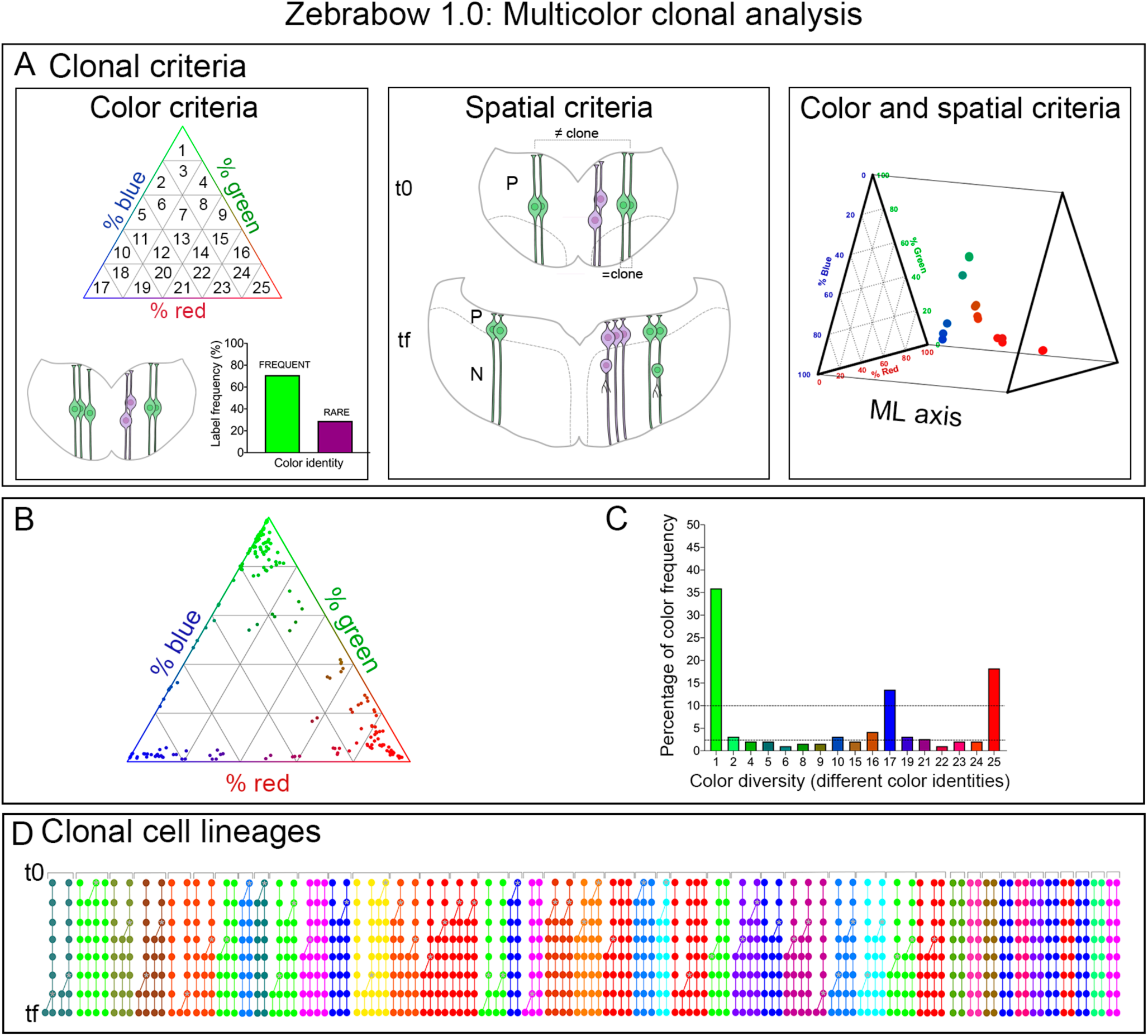
Multicolor clonal tracking using the zebrabow1.0 system. (A) Criteria based on cell color and position used for clonal identification. For the color criterion (left panel), we analyzed the frequencies of color label combinations expressed by boundary cells and used the rarest colors to establish the size and fate criteria at t0. Illustrative example of clones displaying frequent and rare colors. The spatial criterion (middle panel) considered whether cells were in the progenitor (ventricular) at t0; and whether cells displaying the same color were in close contact. We only followed clones with all cells in the progenitor domain at t0 and with cells in close contact displaying the same color. Both color and spatial criteria were used for the clonal assessment (right panel). 3D ternary plot with the mediolateral (ML) axis of r4/r5 boundary at 32hpf corresponding to (Figure 3b). (B) Ternary plot showing the normalized RGB values (% of each fluorescent protein to the global signal) of 169 labeled cells from 4 different embryos (e1–e4) at t0 (32hpf). (C) Histogram displaying the frequency of the 25 color subdivisions defined in (A). The lower black dashed line indicates the 2.5% threshold under which colors were defined as rare. The upper black dashed line indicates the threshold of 10% above which labels were considered frequent. (D) Lineage trees of all the boundary cell clones tracked from 32hpf (t0) to 45hpf (tf) (n=44 clones, N=4 embryos; 90 cells at t0 and 138 cells at tf). Dots represent cells, lines connect the cells from the same track, and branch points indicate cell divisions. Cell lineages are color-coded according to the clone of origin. Time is represented vertically, with t0 at the top and tf at the bottom.

**Figure EV5.**
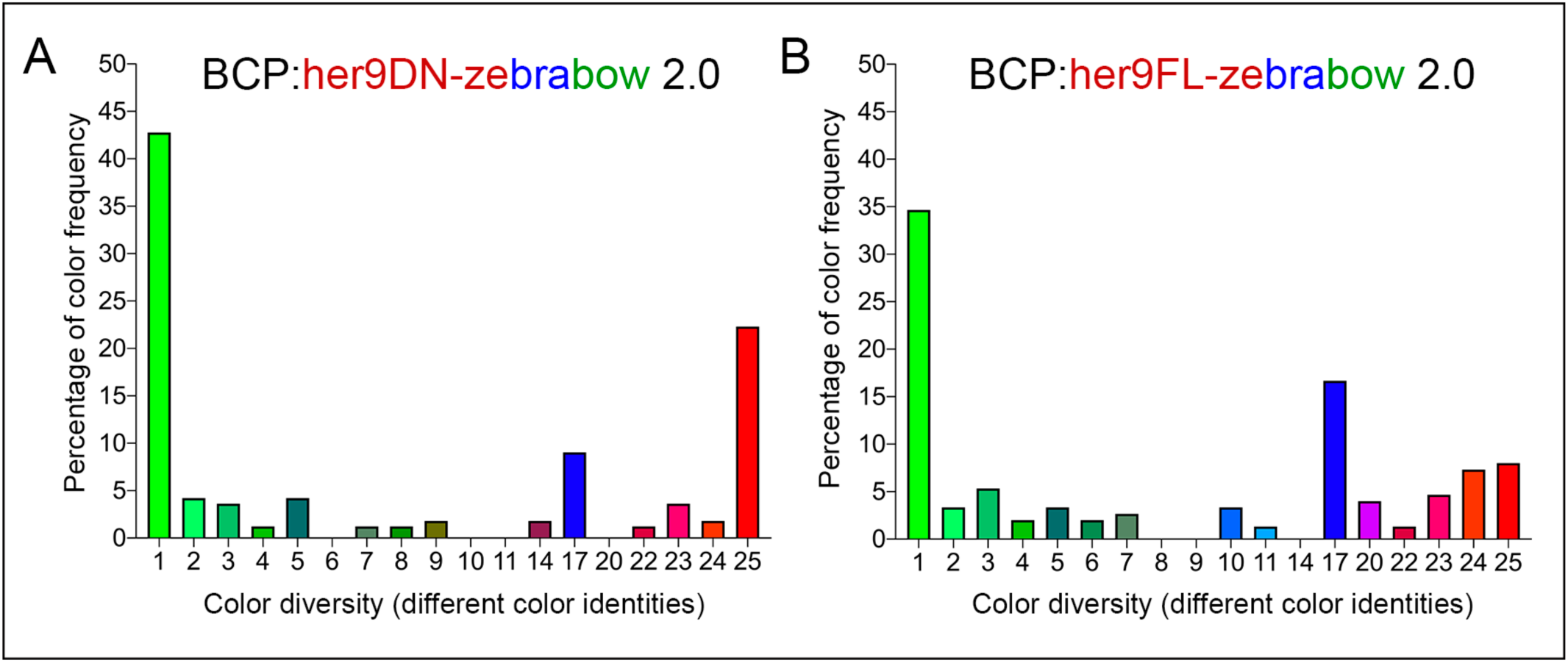
Her9 functional multicolor clonal analysis using the zebrabow 2.0 system. (A–B) Histograms displaying the frequencies of color labels observed at 36hpf in BCP:her9DN-zebrabow2.0 and BCP:her9FL-zebrabow2.0 embryos, respectively. Y-axes represent the percentage of cells found in each of the 25 color subdivisions of the normalized RGB space (only expressed color subdivisions were plotted).

**Figure EV6.**
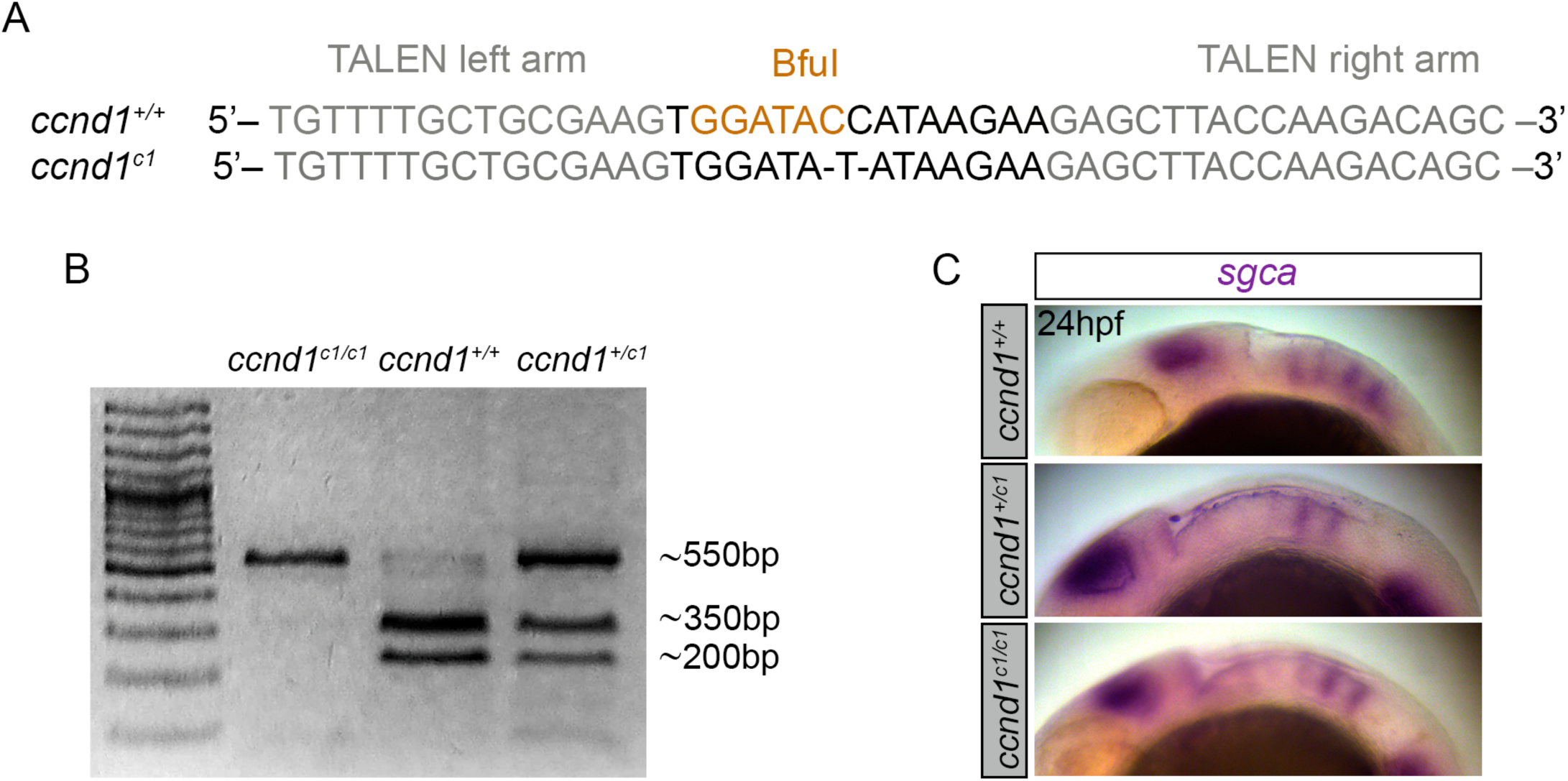
Genomic edition of *ccnd1* gene by TALEN technology. (A) Alignment of the *ccnd1^c1^* mutant allele sequence with the wild-type allele (*ccnd1^+/+^*) showing: i) the deleted nucleotides, and ii) the TALEN target sites in the *ccnd1* locus; the left and right arms (grey) of these sites are separated by a spacer (black) including the restriction site used for screening (orange). (B) Agarose gel with the bands resulting from restriction from embryos carrying the wild-type or the mutated allele. The size of the different obtained fragments is indicated. The *ccnd1^c1^* allele generates a truncated cyclinD1 protein of 12AA. (C) *In situ* hybridization for the boundary marker *sgca* in wild-type (*ccnd1^+/+^*), heterozygous (*ccnd1^+/c1^*), or homozygous (*ccnd1^c1/c1^*) embryos at 24hpf. Note that the mutant *ccnd1^c1/c1^* showed lower *sgca* expression in the most anterior hindbrain boundaries.

**Table EV1.**
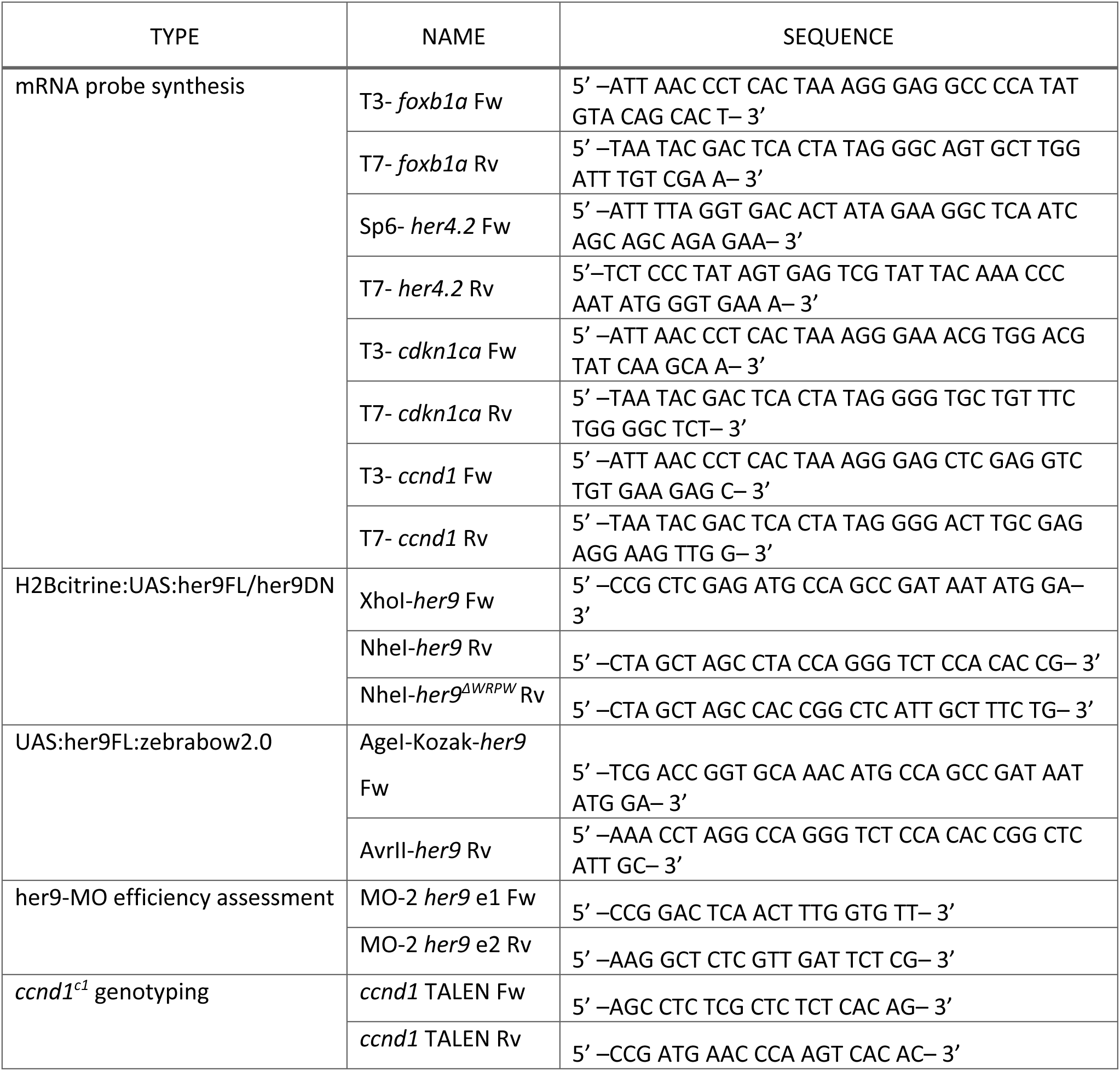

## Notes

### Competing Interest Statement

The authors have declared no competing interest.

## BIBLIOGRAPHY

Amoyel M, Cheng Y-C, Jiang Y-J & Wilkinson DG (2005) Wnt1 regulates neurogenesis and mediates lateral inhibition of boundary cell specification in the zebrafish hindbrain. Development 132: 775–785

Bae Y-K, Shimizu T & Hibi M (2005) Patterning of proneuronal and inter-proneuronal domains by hairy- and enhancer of split-related genes in zebrafish neuroectoderm. Development 132: 1375–1385

Baek JH, Hatakeyama J, Sakamoto S, Ohtsuka T & Kageyama R (2006) Persistent and high levels of Hes1 expression regulate boundary formation in the developing central nervous system. Development 133: 2467–2476

Belmonte-Mateos C, Meister L & Pujades C (2023) Hindbrain rhombomere centers harbor a heterogenous population of dividing progenitors which rely on Notch signaling. Front Cell Dev Biol 11: 1268631

Cau E, Gradwohl G, Casarosa S, Kageyama R & Guillemot F (2000) Hes genes regulate sequential stages of neurogenesis in the olfactory epithelium. Development 127: 2323– 2332

Cermak T, Doyle EL, Christian M, Wang L, Zhang Y, Schmidt C, Baller JA, Somia NV, Bogdanove AJ & Voytas DF (2011) Efficient design and assembly of custom TALEN and other TAL effector-based constructs for DNA targeting. Nucleic Acids Research 39: e82– e82

Chapouton P, Adolf B, Leucht C, Tannhäuser B, Ryu S, Driever W & Bally-Cuif L (2006) her5 expression reveals a pool of neural stem cells in the adult zebrafish midbrain. Development 133: 4293–4303

Cheng Y-C, Amoyel M, Qiu X, Jiang Y-J, Xu Q & Wilkinson DG (2004) Notch activation regulates the segregation and differentiation of rhombomere boundary cells in the zebrafish hindbrain. Developmental Cell 6: 539–550

Clark BS, Cui S, Miesfeld JB, Klezovitch O, Vasioukhin V & Link BA (2012) Loss of Llgl1 in retinal neuroepithelia reveals links between apical domain size, Notch activity and neurogenesis. Development 139: 1599–1610

Dirian L, Galant S, Coolen M, Chen W, Bedu S, Houart C, Bally-Cuif L & Foucher I (2014) Spatial regionalization and heterochrony in the formation of adult pallial neural stem cells. Developmental Cell 30: 123–136

Distel M, Wullimann MF & Köster RW (2009) Optimized Gal4 genetics for permanent gene expression mapping in zebrafish. Proceedings of the National Academy of Sciences of the United States of America 106: 13365–13370

Dong Z, Yang N, Yeo S-Y, Chitnis A & Guo S (2012) Intralineage Directional Notch Signaling Regulates Self-Renewal and Differentiation of Asymmetrically Dividing Radial Glia. Neuron 74: 65–78

Dyballa S, Savy T, Germann P, Mikula K, Remesikova M, Špir R, Zecca A, Peyriéras N & Pujades C (2017) Distribution of neurosensory progenitor pools during inner ear morphogenesis unveiled by cell lineage reconstruction. eLife 6: 951

Engel-Pizcueta C & Pujades C (2021) Interplay Between Notch and YAP/TAZ Pathways in the Regulation of Cell Fate During Embryo Development. Frontiers Cell Dev Biology 9: 711531

Fisher AL, Ohsako S & Caudy M (1996) The WRPW motif of the hairy-related basic helix-loop-helix repressor proteins acts as a 4-amino-acid transcription repression and protein-protein interaction domain. Mol Cell Biol 16: 2670–7

Geling A, Itoh M, Tallafuß A, Chapouton P, Tannhäuser B, Kuwada JY, Chitnis AB & Bally-Cuif L (2003) bHLH transcription factor Her5 links patterning to regional inhibition of neurogenesis at the midbrain-hindbrain boundary. Development 130: 1591–1604

Georgia S, Soliz R, Li M, Zhang P & Bhushan A (2006) p57 and Hes1 coordinate cell cycle exit with self-renewal of pancreatic progenitors. Dev Biol 298: 22–31

Gerety SS & Wilkinson DG (2011) Morpholino artifacts provide pitfalls and reveal a novel role for pro-apoptotic genes in hindbrain boundary development. Developmental Biology 350: 279–289

Goedhart J, Stetten D von, Noirclerc-Savoye M, Lelimousin M, Joosen L, Hink MA, Weeren L van, Gadella TWJ & Royant A (2012) Structure-guided evolution of cyan fluorescent proteins towards a quantum yield of 93%. Nat Commun 3: 751

Gonzalez-Quevedo R, Lee Y, Poss KD & Wilkinson DG (2010) Neuronal regulation of the spatial patterning of neurogenesis. Developmental Cell 18: 136–147

Grison A & Atanasoski S (2020) Cyclins, Cyclin-Dependent Kinases, and Cyclin-Dependent Kinase Inhibitors in the Mouse Nervous System. Mol Neurobiol 57: 3206–3218

Guillemot F (2007) Spatial and temporal specification of neural fates by transcription factor codes. Development 134: 3771–3780

Guthrie S & Lumsden A (1991) Formation and regeneration of rhombomere boundaries in the developing chick hindbrain. Development 112: 221–229

Haddon C, Jiang YJ, Smithers L & Lewis J (1998) Delta-Notch signalling and the patterning of sensory cell differentiation in the zebrafish ear: evidence from the mind bomb mutant. Development 125: 4637–4644

Hatakeyama J, Bessho Y, Katoh K, Ookawara S, Fujioka M, Guillemot F & Kageyama R (2004) Hes genes regulate size, shape and histogenesis of the nervous system by control of the timing of neural stem cell differentiation. Development 131: 5539–5550

Hatakeyama J & Kageyama R (2006) Notch1 Expression Is Spatiotemporally Correlated with Neurogenesis and Negatively Regulated by Notch1-Independent Hes Genes in the Developing Nervous System. Cereb Cortex 16: i132–i137

He J, Zhang G, Almeida AD, Cayouette M, Simons BD & Harris WA (2012) How Variable Clones Build an Invariant Retina. Neuron 75: 786–798

Hevia CF, Engel-Pizcueta C, Udina F & Pujades C (2022) The neurogenic fate of the hindbrain boundaries relies on Notch3-dependent asymmetric cell divisions. Cell Reports 39: 110915

Hirata H, Tomita K, Bessho Y & Kageyama R (2001) Hes1 and Hes3 regulate maintenance of the isthmic organizer and development of the mid/hindbrain. EMBO J 20: 4454–4466

Ishibashi M, Ang SL, Shiota K, Nakanishi S, Kageyama R & Guillemot F (1995) Targeted disruption of mammalian hairy and Enhancer of split homolog-1 (HES-1) leads to up-regulation of neural helix-loop-helix factors, premature neurogenesis, and severe neural tube defects. Genes Dev 9: 3136–3148

Itoh M & Chitnis AB (2001) Expression of proneural and neurogenic genes in the zebrafish lateral line primordium correlates with selection of hair cell fate in neuromasts. MECHANISMS OF DEVELOPMENT 102: 263–266

Kageyama R, Ohtsuka T, Shimojo H & Imayoshi I (2008) Dynamic Notch signaling in neural progenitor cells and a revised view of lateral inhibition. Nature Neuroscience 11: 1247– 1251

Kiecker C & Lumsden A (2005) Compartments and their boundaries in vertebrate brain development. Nature Reviews Neuroscience 6: 553–564

Kim JH, Lee S-R, Li L-H, Park H-J, Park J-H, Lee KY, Kim M-K, Shin BA & Choi S-Y (2011) High Cleavage Efficiency of a 2A Peptide Derived from Porcine Teschovirus-1 in Human Cell Lines, Zebrafish and Mice. PLoS ONE 6: e18556

Kressmann S, Campos C, Castanon I, Fürthauer M & González-Gaitán M (2015) Directional Notch trafficking in Sara endosomes during asymmetric cell division in the spinal cord. Nat Cell Biol 17: 333–339

Labalette C, Bouchoucha YX, Wassef MA, Gongal PA, Men JL, Becker T, Gilardi-Hebenstreit P & Charnay P (2011) Hindbrain patterning requires fine-tuning of early krox20 transcription by Sprouty 4. Development 138: 317–326

Langheinrich U, Hennen E, Stott G & Vacun G (2002) Zebrafish as a model organism for the identification and characterization of drugs and genes affecting p53 signaling. Current Biology 12: 2023–2028

Letelier J, Terriente J, Belzunce I, Voltes A, Undurraga CA, Polvillo R, Devos L, Tena JJ, Maeso I, Rétaux S, et al (2018) Evolutionary emergence of the rac3b/rfng/sgca regulatory cluster refined mechanisms for hindbrain boundaries formation. Proceedings of the National Academy of Sciences of the United States of America 115

Leve C, Gajewski M, Rohr K & Tautz D (2001) Homologues of c-hairy1 (her9) and lunatic fringe in zebrafish are expressed in the developing central nervous system, but not in the presomitic mesoderm. Dev Genes Evol 211: 493–500

Loulier K, Barry R, Mahou P, Franc YL, Supatto W, Matho KS, Ieng S, Fouquet S, Dupin E, Benosman R, et al (2014) Multiplex cell and lineage tracking with combinatorial labels. Neuron 81: 505–520

Lumsden A & Keynes R (1989) Segmental patterns of neuronal development in the chick hindbrain. Nature 337: 424–428

Maeda Y, Isomura A, Masaki T & Kageyama R (2023) Differential cell-cycle control by oscillatory versus sustained Hes1 expression via p21. Cell Reports 42: 112520

Mamidi A, Prawiro C, Seymour PA, Lichtenberg KH de, Jackson A, Serup P & Semb H (2018) Mechanosignalling via integrins directs fate decisions of pancreatic progenitors. Nature 564: 114–118

März M, Chapouton P, Diotel N, Vaillant C, Hesl B, Takamiya M, Lam CS, Kah O, Bally-Cuif L & Strähle U (2010) Heterogeneity in progenitor cell subtypes in the ventricular zone of the zebrafish adult telencephalon. Glia 58: 870–888

McLarren KW, Theriault FM & Stifani S (2001) Association with the Nuclear Matrix and Interaction with Groucho and RUNX Proteins Regulate the Transcription Repression Activity of the Basic Helix Loop Helix Factor Hes1*. J Biol Chem 276: 1578–1584

Miesfeld JB & Link BA (2014) Establishment of transgenic lines to monitor and manipulate Yap/Taz-Tead activity in zebrafish reveals both evolutionarily conserved and divergent functions of the Hippo pathway. MECHANISMS OF DEVELOPMENT 133: 177–188

Mizutani K, Yoon K, Dang L, Tokunaga A & Gaiano N (2007) Differential Notch signalling distinguishes neural stem cells from intermediate progenitors. Nature 449: 351–355

Moens CB & Prince VE (2002) Constructing the hindbrain: insights from the zebrafish. Developmental Dynamics 224: 1–17

Moens CB, Yan Y-L, Appel B, Force AG & Kimmel CB (1996) valentino: a zebrafish gene required for normal hindbrain segmentation. Development 122: 3981–3990

Monahan P, Rybak S & Raetzman LT (2009) The Notch Target Gene Hes1 Regulates Cell Cycle Inhibitor Expression in the Developing Pituitary. Endocrinology 150: 4386–4394

Nerli E, Rocha-Martins M & Norden C (2020) Asymmetric neurogenic commitment of retinal progenitors involves Notch through the endocytic pathway. eLife 9

Nikolaou N, Watanabe-Asaka T, Gerety S, Distel M, Köster RW & Wilkinson DG (2009) Lunatic fringe promotes the lateral inhibition of neurogenesis. Development 136: 2523– 2533

Ninkovic J, Tallafuss A, Leucht C, Topczewski J, Tannhäuser B, Solnica-Krezel L & Bally-Cuif L (2004) Inhibition of neurogenesis at the zebrafish midbrain-hindbrain boundary by the combined and dose-dependent activity of a new hairy/E(spl)gene pair. Development 132: 75–88

Ohtsuka T, Ishibashi M, Gradwohl G, Nakanishi S, Guillemot F & Kageyama R (1999) Hes1 and Hes5 as Notch effectors in mammalian neuronal differentiation. EMBO J 18: 2196– 2207

Ohtsuka T & Kageyama R (2021) Hes1 overexpression leads to expansion of embryonic neural stem cell pool and stem cell reservoir in the postnatal brain. Development 148

Pan YA, Freundlich T, Weissman TA, Schoppik D, Wang XC, Zimmerman S, Ciruna B, Sanes JR, Lichtman JW & Schier AF (2013) Zebrabow: multispectral cell labeling for cell tracing and lineage analysis in zebrafish. Development 140: 2835–2846

Park HC, Kim CH, Bae YK, Yeo SY, Kim SH, Hong SK, Shin J, Yoo KW, Hibi M, Hirano T, et al (2000) Analysis of upstream elements in the HuC promoter leads to the establishment of transgenic zebrafish with fluorescent neurons. Developmental Biology 227: 279–293

Park S-H, Yeo S-Y, Yoo K-W, Hong S-K, Lee S, Rhee M, Chitnis AB & Kim C-H (2003) Zath3, a neural basic helix-loop-helix gene, regulates early neurogenesis in the zebrafish. BIOCHEMICAL AND BIOPHYSICAL RESEARCH COMMUNICATIONS 308: 184–190

Peretz Y, Eren N, Kohl A, Hen G, Yaniv K, Weisinger K, Cinnamon Y & Sela-Donenfeld D (2016) A new role of hindbrain boundaries as pools of neural stem/progenitor cells regulated by Sox2. BMC biology 14: 57

Pujades C (2020) The multiple functions of hindbrain boundary cells: Tinkering boundaries? Seminars in Cell and Developmental Biology 107: 179–189

Revenu C, Revenu C, Streichan S, Streichan S, Donà E, Dona E, Lecaudey V, Lecaudey V, Hufnagel L, Hufnagel L, et al (2014) Quantitative cell polarity imaging defines leader-to-follower transitions during collective migration and the key role of microtubule-dependent adherens junction formation. Development 141: 1282–1291

Shaner NC, Campbell RE, Steinbach PA, Giepmans BNG, Palmer AE & Tsien RY (2004) Improved monomeric red, orange and yellow fluorescent proteins derived from Discosoma sp. red fluorescent protein. Nat Biotechnol 22: 1567–1572

Shcherbakova DM & Verkhusha VV (2013) Near-infrared fluorescent proteins for multicolor in vivo imaging. Nat methods 10: 751–4

Shimojo H, Ohtsuka T & Kageyama R (2008) Oscillations in notch signaling regulate maintenance of neural progenitors. Neuron 58: 52–64

Sigloch C, Spitz D & Driever W (2023) A network of Notch-dependent and -independent her genes controls neural stem and progenitor cells in the zebrafish thalamic proliferation zone. Development 150: dev201301

Stigloher C, Chapouton P, Adolf B & Bally-Cuif L (2008) Identification of neural progenitor pools by E(Spl) factors in the embryonic and adult brain. Brain Research Bulletin 75: 266– 273

Sueda R, Imayoshi I, Harima Y & Kageyama R (2019) High Hes1 expression and resultant Ascl1 suppression regulate quiescent vs. active neural stem cells in the adult mouse brain. Genes & Development 33: 511–523

Thisse C & Thisse B (2008) High-resolution in situ hybridization to whole-mount zebrafish embryos. Nature Protocols 3: 59–69

Voltes A, Hevia CF, Engel-Pizcueta C, Dingare C, Calzolari S, Terriente J, Norden C, Lecaudey V & Pujades C (2019) Yap/Taz-TEAD activity links mechanical cues to progenitor cell behavior during zebrafish hindbrain segmentation. Development 146

Wilkinson MF & Shyu A (2001) Multifunctional regulatory proteins that control gene expression in both the nucleus and the cytoplasm. BioEssays 23: 775–787

Wolff C, Tinevez J-Y, Pietzsch T, Stamataki E, Harich B, Guignard L, Preibisch S, Shorte S, Keller PJ, Tomancak P, et al (2018) Multi-view light-sheet imaging and tracking with the MaMuT software reveals the cell lineage of a direct developing arthropod limb. eLife 7: 375

Zacharias DA, Violin JD, Newton AC & Tsien RY (2002) Partitioning of Lipid-Modified Monomeric GFPs into Membrane Microdomains of Live Cells. Science 296: 913–916

Zalc A, Hayashi S, Auradé F, Bröhl D, Chang T, Mademtzoglou D, Mourikis P, Yao Z, Cao Y, Birchmeier C, et al (2014) Antagonistic regulation of p57kip2 by Hes/Hey downstream of Notch signaling and muscle regulatory factors regulates skeletal muscle growth arrest.

